# Bulb growth potential is independent of leaf longevity for the spring ephemeral *Erythronium americanum* Ker-Gawl

**DOI:** 10.1101/2022.07.19.500717

**Authors:** Hugo Bertrand, Line Lapointe

## Abstract

Growth in most spring ephemerals is decreased under warmer temperatures. Although photosynthetic activities are improved at warmer temperatures, leaves senesce earlier, which prevents the bulb from reaching a larger size. A longer leaf life duration during a warm spring, therefore, may improve bulb mass. We tested this hypothesis by modulating leaf life span of *Erythronium americanum* through the application of Promalin® (PRO; cytokinins and gibberellins) that prolonged, or silver thiosulphate (STS) that reduced leaf duration. Gas exchange and chlorophyll fluorescence were measured along with leaf and bulb carbohydrate concentrations. Plants were also pulse labelled with ^13^CO_2_ to monitor sugar transport to the bulb. Lower photosynthetic rates and shorter leaf life span of STS plants reduced the quantity of C that they assimilated during the season, resulting in a smaller bulb compared to Control plants. PRO plants maintained their photosynthetic rates for a longer period than Control plants, yet final bulb biomass did not differ between them. We conclude that seasonal growth for *E. americanum* is not limited by leaf life duration under warm growing conditions, but rather by limited sink growth capacity. Under global warming, spring geophytes might be at risk of being reduced in size and eventually, reproducing less frequently.

**Highlight:** Warmer springs negatively affect trout lily growth and delaying leaf senescence in this spring ephemeral does not translate into a larger bulb if temperatures remain high during springtime.

## Introduction

Temperate forest spring ephemerals, such as yellow trout lily (*Erythronium americanum* Ker-Gawler), have evolved to take advantage of the high irradiance conditions between snowmelt and overstory canopy closure (Muller, 1978). Thus, they only have 40-60 days to complete their epigeous growth. Since *E. americanum* can be characterized as a true “sun” plant, photosynthetic rates are high during this period (Sparling, 1967; Taylor and Pearcy, 1976; Lapointe, 2001; Recchia *et al*., 2017). These high photosynthetic rates allow fast growth and a high starch accumulation rate in the bulb, which will be a major determinant of subsequent hypogeous growth during winter and, eventually, of reproduction and survival.

The timing of canopy closure usually determines leaf longevity of spring ephemerals, such as ramps (*Allium tricoccum* Aiton) and yellow star-of-Bethlehem (*Gagea lutea* [L.] Ker Gawler), and influences their annual growth (Kim *et al*., 2015; Dion *et al*., 2017). Light reduction caused by tree leaf-out is an important cue for senescence of spring ephemerals. Yet, canopy closure is not the only abiotic factor affecting leaf longevity of spring ephemerals, given that temperature can also modulate the timing of senescence and seasonal growth (Yoshie, 2008; Bernatchez and Lapointe, 2012; Sunmonu and Kudo, 2015). Warmer temperatures usually hasten leaf senescence and negatively affect growth. Leaf emergence of *E. americanum* can also be hastened under earlier snowmelt (Tessier, 2019). Climate change affects both snow cover duration and average spring temperature in Quebec, Canada (Brown, 2010; Vincent *et al*., 2012). Canopy closure occurs earlier with climate change, but the phenology of tree leaf-out varies across species and depends upon a combination of factors such as winter chilling, spring temperature and photoperiod (Polgar and Primack, 2011). Nevertheless, increasing spring temperature has been shown to decrease leaf life span of most spring ephemerals, even if they emerge earlier (Augspurger and Zaya, 2020).

Spring ephemerals, as well as many other geophytes, are unique in that they produce larger perennial organs at cooler (8-12°C) than at warmer temperatures (Badri *et al*., 2007; Gandin *et al*., 2011b; Bernatchez and Lapointe, 2012). Sustained source-sink imbalance has been proposed as the mechanism by which higher temperature negatively affects growth in *E. americanum* (Gandin *et al*., 2011b). Enhanced photosynthetic rates that is caused by warmer temperatures appears to exceed sink demand, giving rise to feedback inhibition of photosynthesis and, eventually to leaf senescence. Indeed, it is well established that photosynthetic rates are regulated by sink tissues to ensure a balance between source and sink strengths (Paul and Foyer, 2001). Under limited sink demand, a steeper and earlier decrease in photosynthetic rate in combination with early leaf senescence appears to prevent bulb and bulb cells from achieving the sizes reached at lower temperatures (Gandin *et al*., 2011b). However, such reductions in cell growth cannot fully be explained by decreasing photosynthetic rates, given that enhancing photosynthetic rates through CO_2_ enrichment or increased irradiance does not improve bulb growth (Gandin *et al*., 2009; Gandin *et al*., 2011a). Further, it has been shown that the bulb can cope with the daily excess of carbon that is assimilated under these high photosynthetic rates by stimulating the alternative respiratory pathway (Gandin *et al*. 2009; Dong *et al*., 2018) to restore source-sink balance. Shortened leaf longevity that is caused by the early and sustained source-sink imbalance appears to be the most likely explanation as to why bulbs reach a smaller size at warmer temperatures. Marcelis (1996) suggested that sink strength is better defined as the growth potential of an organ rather than as its achieved growth. Assuming that the intrinsic growth potential of the bulb is not influenced by temperature means that cell size can be increased under warm temperatures; if we increase the duration of the source-sink interaction in *E. americanum*, then the carbon assimilated during the additional days could be used to support further bulb growth.

The aim of this study is to investigate the effect of leaf longevity on the growth of *E. americanum* at warm temperatures. Leaf life span was modulated by spraying Promalin® (PRO, a mixture of gibberellins 4 and 7, and benzyladenine) or silver thiosulphate (STS) on leaves to delay or hasten senescence, respectively. Leaf photosynthetic rates were monitored regularly, which allowed us to calculate total carbon assimilation under different leaf longevities. Leaf and bulb respiratory rates, together with leaf chlorophyll fluorescence, were measured to identify potential mechanisms that may be induced to cope with additional carbon assimilated under a longer growing season. Pulse labelling with ^13^C allowed us to quantify sugar translocation. When coupled with sugar quantification in the leaf and bulb, this estimate of C translocation helped us to pinpoint the presence of a potential sugar feedback from the bulb. This study brings a better understanding of the effects of source longevity in the case of sink-limited growth. More specifically, it allowed us to verify whether artificially lengthening the source-sink interaction is sufficient to exploit the sink growth potential fully when a strong imbalance reduces leaf life span.

## Material & methods

### Plant Material and Growth Conditions

In September 2018, 500 bulbs of *Erythronium americanum* were collected near Sainte- Brigitte-de-Laval (46.93° N, -71.16° W). The following day, bulbs were planted at a depth of 6.5 cm in 0.46 L pots (9 cm deep with a radius of 5 cm at the top and 3.5 cm at the bottom) filled with wet Turface (Profile products LLC, Buffalo Grove, IL, USA). The 400 bulbs retained for the experiment had a diameter between 4.6 mm and 10 mm, and a fresh mass ranging from 0.106 g to 1.015 g. Pots were transferred to a growth cabinet at 20 °C. Every two weeks, temperature was decreased by 4 °C, until it reached 4 °C on 1 November 2018. The pots were then transferred to a cold room for 4.5 months and watered when needed to maintain substrate moisture content. Pots were transferred in mid-March into growth chambers set at 8 °C for a few days (Conviron Inc., Winnipeg, MB, Canada). Plants were then randomly assigned to one of three treatments, in two growth chambers. Growing conditions were identical in both chambers with an irradiance of 300 µmol photons m^-2^ s^-1^, air temperature of 18 °C (day) and 14 °C (night), a photoperiod of 14 h and a relative humidity of 75%. Plants were watered daily and fertilized once a week with 150 mL of 10% Hoagland solution for optimal growth (Lapointe and Lerat, 2006).

Plants from the first treatment were sprayed twice with a solution of 2 mM STS, which also contained 0.1 % v/v Tween-20 (Cameron and Reid, 1981); those in the second treatment were also sprayed twice with a diluted solution of PRO (Valent BioSciences Corporation, Libertyville, IL) containing 10 µM 6-benzyladenine and 10 µM gibberellin 4 and 7 (Gonzalez-Santos *et al*., 2009). The third final treatment was the Control (C), in which plants were sprayed twice with 0.1% v/v Tween-20. Treatments were applied to leaves 7 days after full expansion, then reapplied 5 days later, except for the STS treatment. In this latter case, the second foliar application was applied 3 days after the first to avoid spraying once leaf senescence had begun. Treatments were selected based on preliminary trials testing the effect of two concentrations each of aminooxyacetic acid (AOA), STS, kinetin, and PRO on leaf longevity (Supplementary data Fig. S1). Surprisingly, both ethylene synthesis inhibitors (aminooxyacetic acid) and ethylene action inhibitors (STS) reduced leaf life span of *E. americanum* in those prior trials. Treatments and concentrations showing greatest effects on leaf longevity were selected for this study.

### Gas Exchange and Chlorophyll a Fluorescence

Leaf gas exchange and chlorophyll a fluorescence were measured on three plants per treatment per growth chamber, for a total of 18 plants. Measurements were performed every two days from complete leaf unfolding to the beginning of leaf senescence, and daily from the beginning of leaf senescence up to 50% (area-based) senescence. Net photosynthetic rates were recorded using a LI-6400XT (LI-COR Biosciences, Lincoln, NE, USA) at an irradiance of 300 µmol photons m^-2^ s^-1^, 500 ppm CO_2_ (corresponding to the mean ambient CO_2_ concentration in and around the growth chambers) and a flow rate of 200 µmol s^-1^. Chlorophyll a fluorescence was recorded at the same time with a LI- COR6400-40 leaf chamber fluorometer. At growth irradiance, a 0.8 s flash of 8000 µmol photon m^-2^ s^-1^ was applied to measure Fm’; then, a far-red pulse of 5 s, overlapping with 5 s without actinic light, which started 1 s after the beginning of the far-red pulse, was applied to measure Fo’. Plants were subsequently placed in total darkness for 40 min to measure leaf respiratory rates and dark fluorescence. Fo and Fm were recorded respectively before and after a 0.8 s flash of 8000 µmol photon m^-2^ s^-1^. Daily carbon assimilation (mg C cm^-2^ day^-1^) was calculated by extending leaf photosynthetic rates to the entire photoperiod, and by subtracting respiration that occurred during the night. We estimated assimilation between measurement days by averaging the previous and following daily assimilation. Total C assimilation over the course of the season was obtained by summing daily assimilation for each of the 18 plants. Once senescence became apparent, daily carbon assimilation per plant was corrected for the proportion that remained green on that date.

Bulb respiratory rates were measured on 4 plants per treatment the day following each foliar application of treatments, then every 3 days following the second foliar application until the beginning of leaf senescence. Thereafter, measurements were taken at 5, 25, 50 and 100% of leaf yellowing. Bulb respiratory rates were measured using an ADC-LCA4 gas exchange measurement system (ADC Bioscientific, Hoddensdon, UK), at a flow of 200 µmol s^-1^. We used the ADC-LCA4 to measure bulb respiratory rates instead of the LI- 6400XT since the cuvette with the fluorometer was too thin to contain the bulb; furthermore, interchanging the head of the IRGA requires recalibration between leaf gas exchange and bulb respiratory rates. Plants used for bulb respiration measurements were subsequently flash-frozen in liquid nitrogen and stored at -80 °C, until extraction and quantification of carbohydrates.

### Plant Growth and Leaf Phenology

Maximum leaf area was measured for each plant about 10 days following complete leaf unfolding, between the first and second foliar spraying of STS and PRO treatments. Photographs of the leaves superimposed on a metric grid allowed non-destructive measures of leaf area using ImageJ software (https://imagej.nih.gov/ij/). Each plant that was harvested to measure bulb respiration was subsequently lyophilized and each organ was weighed. Bulbs and leaves were then ground using a MM400 mixer mill (Retsch GmbH, Haan, Germany) and tungsten beads, for 45 s at 30 Hz. To correct for the great variability in autumnal bulb mass and subsequent leaf area that was produced, plant growth was expressed as bulb mass per leaf area. Leaf senescence was assessed visually; a strong correlation between the first visual sign of leaf senescence and biochemical markers of leaf senescence has been reported for this species (Dong, 2020). The number of days from complete leaf unfolding to 5, 25, 50 and 100% yellowing was monitored for each plant.

### 13CO2 Pulse-Chase Labelling

Six plants per treatment per growth chamber were placed in a 100 cm × 30 cm × 50 cm (L × W × H) acrylic box, 8 days after leaf unfolding, then followed by another group at 5% leaf senescence (2 labelling stages). The box was installed in a growth chamber set at the same temperature and light conditions as the other growth chambers. The CO_2_ concentration in the box (measured with a LI-840 infrared gas analyzer; LI-COR Biosciences) was initially decreased to 450 ppm using soda lime. Subsequently, 99% ^13^CO_2_ was injected into the box to reach a concentration of about 580 ppm (∼20% ^13^CO_2_). Water vapour in this closed system was controlled by circulating box air through a column of CaSO_4_ prior to entering the gas analyzer during pulse labelling. The CO_2_ concentration in the box never dropped below 490 ppm (a reduction of less than 15%) during the course of pulse labelling. After an hour of labelling, plants were removed from the box. Four plants per treatment were harvested and frozen in liquid nitrogen immediately after pulse-labelling, while the other plants were moved back to their growth chambers. At the same time, 4 additional unlabelled plants were harvested as Controls for each treatment and labelling stage. Four pulse-labelled plants per treatment were harvested 24 hours after pulse labelling was accomplished and 4 more were harvested 48 hours after pulse labelling. Bulbs and leaves of pulse-labelled and unlabelled plants were later lyophilized, weighed, and ground following the same method used to characterize plant biomass. A 1 mg subsample was used to assess ^13^C content by gas chromatography-mass spectrometry (laboratory of Jean-Éric Tremblay, Centre de recherche Quebec-Ocean, Université Laval). Leaf and bulb ^13^C content of unlabelled plants was used as a baseline of ^13^C content for pulse-labelled plants. Excess ^13^C content of pulse-labelled plants was calculated by subtracting this baseline from ^13^C of pulse-labelled plants, as determined by GC-MS.

### Non-Structural Carbohydrates Quantification and Plant Biomass

Soluble sugars and starch were extracted from plants that were used for bulb respiration measurements, following the protocol of Landhausser et al. (2018). Thirty mg of ground tissues were mixed with 1.5 mL of ethanol 80%, then heated at 90 °C for 10 minutes. A subsample (0.2 mL) of supernatant was set aside for soluble sugar determination. The pellet was washed twice with ethanol then kept under a fume hood overnight to evaporate the remaining ethanol. Starch in the pellet was digested with α-amylase (MilliporeSigma, Oakville, ON, Canada) at 85 °C for 30 minutes. Thereafter, supernatant containing maltose from starch was digested with amyloglucosidase (MilliporeSigma) at 55 °C for another 30 minutes. Glucose from starch was quantified colorimetrically at 415 nm following a reaction with p-hydroxybenzoic acid hydrazide (MilliporeSigma; Blakeney et al., 1980). Reducing sugars from the initial supernatant were quantified using the same method both before and after sucrose digestion with invertase, allowing us to calculate the quantity of sucrose along with that of reducing sugars.

### Statistical Analyses

All statistical analyses were performed using R 3.6.3 (R Core Team, 2020), with growth chamber as a block factor. Leaf phenology, bulb mass at final harvest and total C assimilated over the course of the season were analyzed by ANOVA with treatment as a fixed factor. Carbohydrate concentration in the leaves and bulbs, and bulb respiratory rates were analyzed by ANOVA in a factorial design using treatments and harvests as fixed factors; ^13^C content in the leaf and bulb were analyzed by ANOVA in a factorial design testing treatments, labelling stage and chase period as fixed factors. Leaf net photosynthetic rates, respiratory rates and chlorophyll fluorescence responses were analyzed by repeated- measures ANOVA using the *lme* function from the *nlme* package in R (Pinheiro et al., 2021), with treatments and time as fixed factors, and plants as a random factor (subject factor). The correlation matrix structure that was used for the mixed model was the continuous first-order autoregressive (CorCAR1) for all analyses and was selected by AIC (Akaike Information Criterion) from among other correlation models. Bulb growth rate per leaf area was tested with multiple growth rate models, and the best model was selected by AIC. An additional sums-of-squares term was applied to the selected null model and its grouped counterpart to detect differences among treatments (Ritz and Streibig, 2008). Plants that produced rhizomes were excluded from the analyses, since the effect of rhizome production on leaf longevity and belowground mass accumulation has yet to be studied and may alter the source-sink balance.

## Results

### Plant Phenology and Growth

Treatments were applied 7 and 12 days (7 and 10 days for STS) after complete leaf unfolding and induced an earlier onset of leaf senescence for STS plants (mean ± SE: 13.5 ± 0.4 days) and a delayed onset for PRO plants (23.2 ± 0.8 days) as compared to Control plants (21 ± 0.6 days; *F_2,243_* = 63.8, *P* < 0.001, Fig. 1). Complete leaf senescence occurred at 21.7 ± 0.7 days for STS, 30.2 ± 0.6 days for Control and 36.6 ± 0.6 days for PRO plants (*F_2,135_* = 135, *P* < 0.001). This represents an increase of 6.4 days or 21% of the leaf life span for PRO plants, and a decrease of 8.5 days, or 28%, for STS plants when compared to Control plants. The duration of senescence was longer for PRO plants (13.3 ± 0.8 days) than STS (7.3 ± 0.4 days) and Control plants (7.4 ± 0.4 days; *F_2,135_* = 17.5, *P* < 0.001).

**Fig. 1:**
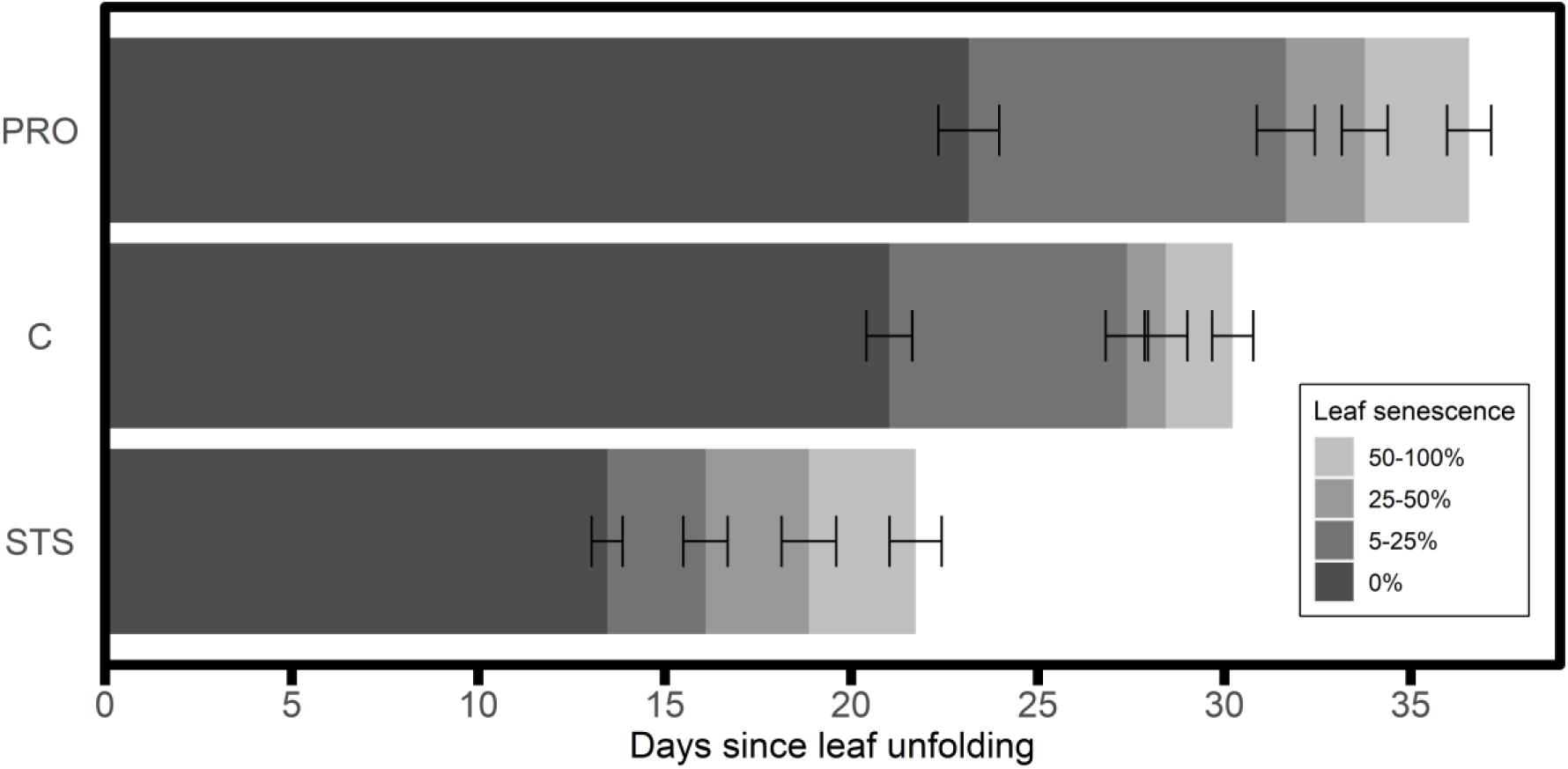
Number of days that elapsed between complete leaf unfolding and 5, 25, 50 or 100% leaf yellowing for *E. americanum* plants treated with Promalin (PRO), silver thiosulphate (STS), or water (C). Bars represent mean duration of the different phenological stages ± 1 standard error.

Reducing leaf life span with STS also decreased bulb dry mass at final harvest (187 ± 11 mg) when compared to Control plants (298 ± 15 mg; *F*_2,135_ = 20.4, *P* < 0.001). This represents a reduction of 26% of bulb mass and is consistent with the decrease in leaf life span (28%). Despite a delay in the senescence process for PRO plants, their bulb dry mass (278 ± 12 mg) was not higher than that of Control plants (298 ± 15 mg). Treatments did not affect leaf area, with a mean leaf area of 8.1 ± 0.3 cm^2^ across treatments (*F*_2,250_ = 0.44, *P* = 0.65). Plant leaf area showed no correlation with the time elapsed between leaf unfolding and either the beginning of senescence or its completion for any of the treatments (Supplementary Table S1). However, plant leaf area of Control plants was strongly correlated with bulb mass at final harvest (*r* = 0.94), even more so than with the autumnal bulb mass (i.e., the fresh mass recorded when bulbs were collected in the forest in the previous autumn; Fig. 2, *r* = 0.53; Supplementary Table S2). The variability in leaf area between individuals lead to great variation in bulb dry mass by the end of the growing season, masking possible differences between treatments (Fig. 3A). To account for the variation in leaf size, bulb mass was also expressed per leaf area, i.e., the maximum leaf area attained by the plant early in the season (Fig. 3B). Final bulb mass that was reached per leaf area was reduced for STS (28.6 ± 1.1 mg cm^-2^), but did not differ between PRO (40.7 ± 1.0 mg cm^-2^) and Control plants (38.6 ± 0.7 mg cm^-2^; *F*_2,131_ = 52.8, *P* < 0.001), thereby confirming the lack of an effect of extended leaf life duration on final bulb mass.

**Fig. 2:**
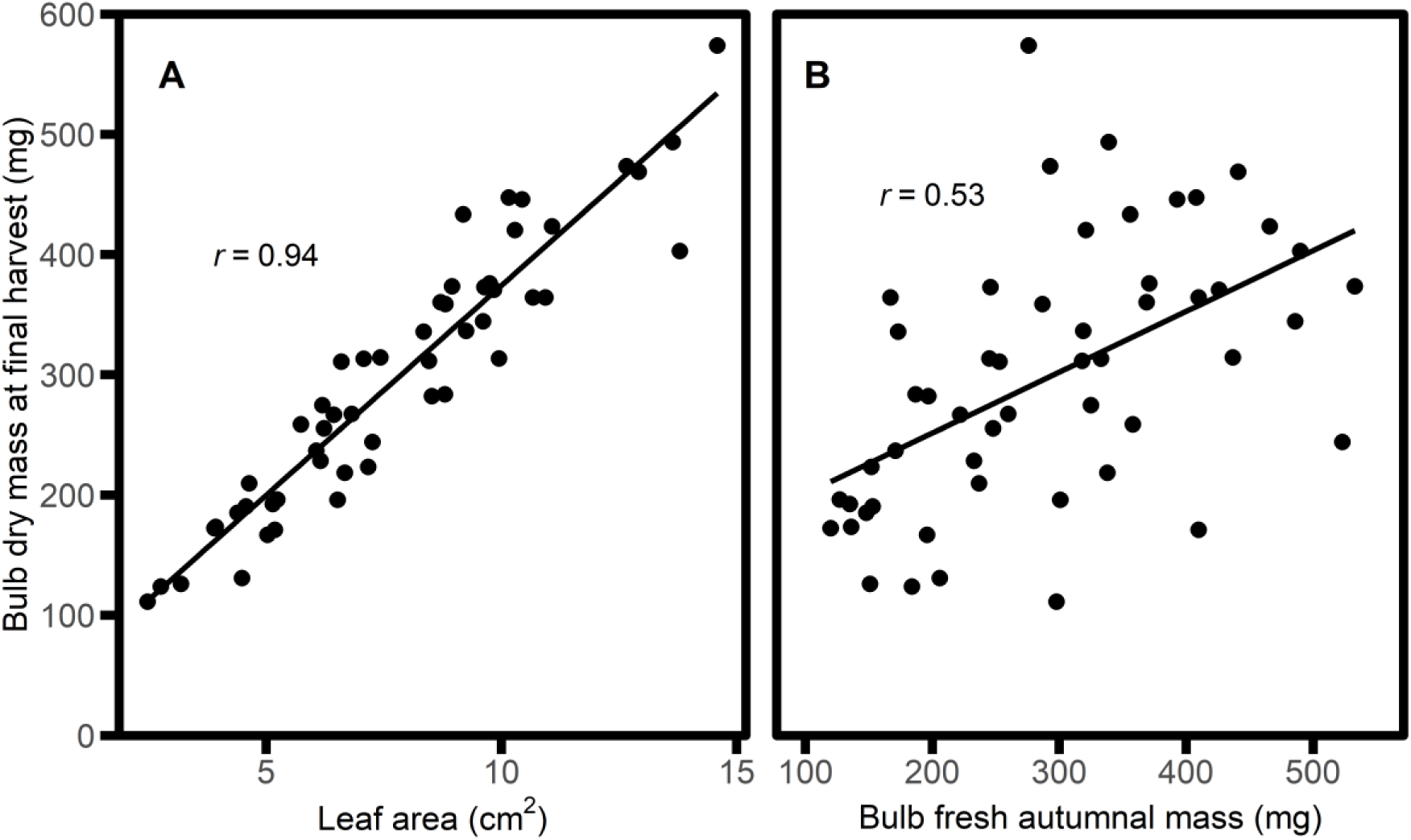
Relationship between either fully expanded leaf area (left) or bulb autumnal fresh mass (right) and bulb dry mass at final harvest for Control plants of *E. americanum*. Each data point represents an individual plant harvested at 100% senescence. Both plots have the same dataset of plants (*n* = 52).

**Fig. 3:**
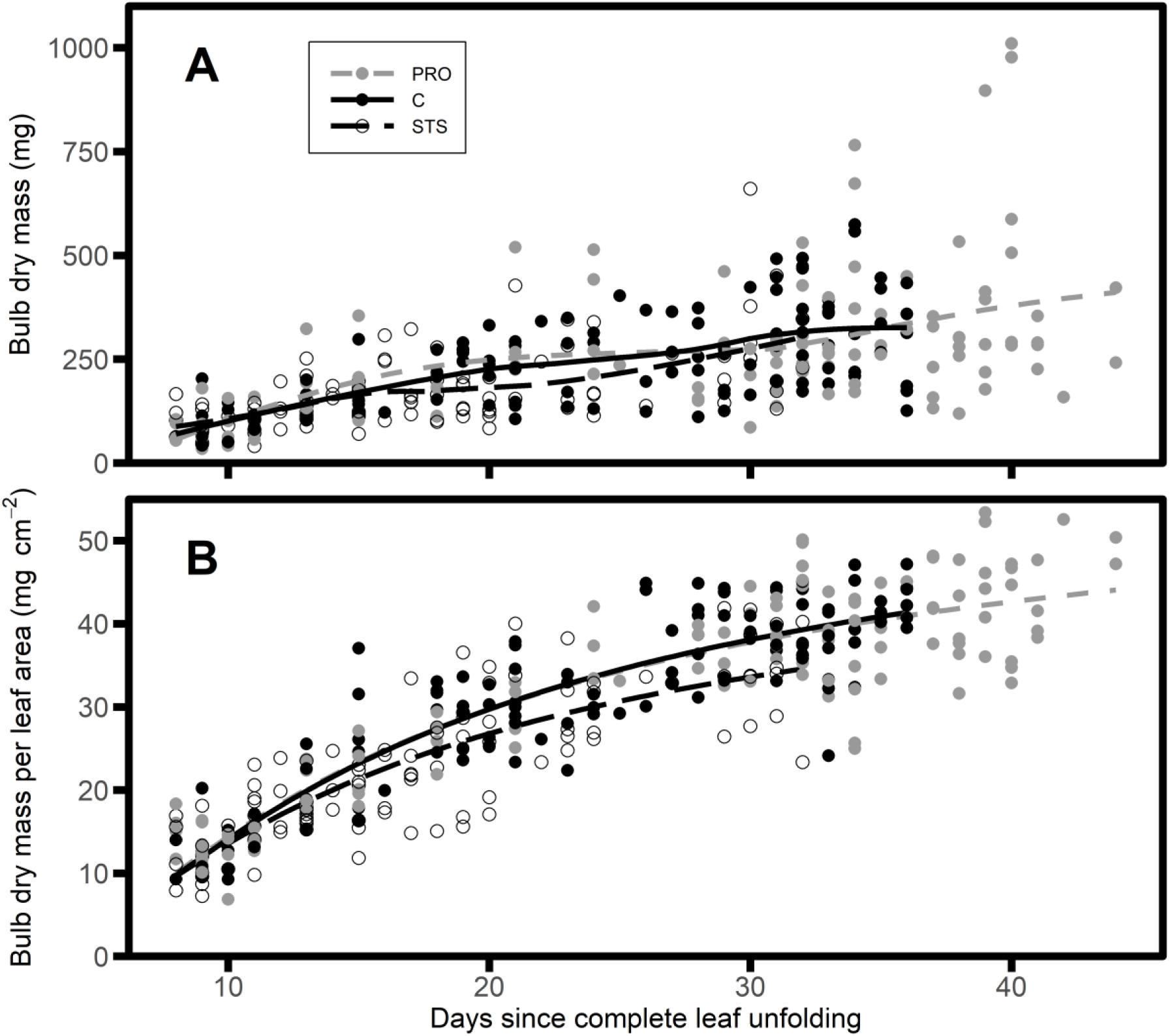
Bulb dry mass (panel A) and bulb dry mass per leaf area (panel B; BLA) increased through time following completed leaf unfolding in *E. americanum* plants. Each data point in panel A represents the bulb mass of an individual plant, and the BLAs of the same plants are presented in panel B, i.e., the bulb biomass at the time of harvest divided by the maximum leaf area reached earlier in the season. The curves on panel A represent LOESS regressions showing the general pattern of the data points, and the three curves on panel B represent the logarithmic reciprocal growth curve for each treatment: foliar application of Promalin (PRO, grey), silver thiosulphate (STS, white) or water (C, black). LOESS: locally estimated scatterplot smoothing (Wickham, 2016).

Bulb dry mass per leaf area, which we determined using plants that were sampled for bulb respiration measurements (BLA, mg cm^-2^), increased through time following a logarithmic reciprocal model, as shown in Fig. 3B. The growth curve for each treatment followed this equation: 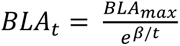, where *t* represents the number of days since complete leaf unfolding; BLA_max_ is the maximum BLA value, and β is a growth parameter (Table 1). What stands out in Fig. 3 is the large difference between the curve for STS plants as compared to those of the two other treatments. Indeed, there was a significant difference between the null model and the grouped model (*F_4,336_* = 5.48; *P* < 0.001), suggesting that at least one of the treatments differed from the others. The STS curve differed from that of the Control plants (*F_2,221_* = 10.0, *P* < 0.001), but PRO and Control plants were not different from one another (*F_2,235_* = 0.11, *P* = 0.9). Both parameters (BLA_max_ and β) of the curve significantly differed between STS and Control plants (*P* < 0.001 for both parameters). This indicates that the reduction of growth in STS plants was not solely explained by a reduction in leaf life span; otherwise, the three curves would have been identical. STS treatment seemed to affect bulb growth in addition to reducing leaf life span. Another striking observation in this figure was the similarity between PRO and Control growth curves. Prolonging leaf life span with the PRO treatment did not seem to affect the growth of the plant, but only lengthened the growing season. The growth rate near the end of the growing season basically plateaued, allowing very little additional growth when lengthening leaf life span.

**Table 1:** Means and standard errors of the parameters of the logarithmic reciprocal growth curve of *E. americanum* following the spraying (foliar application) of Promalin (PRO), silver thiosulphate (STS), or water (C). BLA_t_ is expressed as bulb dry mass per leaf area (mg cm^-2^) on a given day since complete leaf unfolding. BLA_max_ is the asymptotic upper BLA threshold, whereas β is a growth parameter and *t* represents the number of days since complete leaf unfolding.

| Treatments | $\beta$ | $BLA_{\max}$ | Growth curve |
| --- | --- | --- | --- |
| Promalin (PRO) | $14.5 \pm 0.9$ | $61.3 \pm 1.9$ | $BLA_t = \frac{61.3}{e^{14.5/t}}$ |
| Control (C) | $14.9 \pm 0.9$ | $62.6 \pm 2.4$ | $BLA_t = \frac{62.6}{e^{14.9/t}}$ |
| Silver thiosulphate (STS) | $13.6 \pm 1.0$ | $52.9 \pm 3.1$ | $BLA_t = \frac{52.9}{e^{13.6/t}}$ |

### Gas Exchange Measurements

Leaf respiratory rate remained constant throughout the growing season and exhibited no difference among treatments (Table 2). Bulb respiratory rates showed an initial decrease from Day 7 to Day 15 after leaf unfolding for STS and from Day 7 to Day 20 for Control and PRO plants (Fig. 4). Thereafter, bulb respiratory rates exhibited a temporary increase, which corresponded to the point at which leaf senescence began in all three treatments. There was no difference in bulb respiratory rates within each phenological stage among the three treatments (Table 2). Nevertheless, bulbs of PRO plants likely respired more carbon during leaf senescence, since this period lasted longer for PRO plants than for the two other treatment groups. In contrast to the leaf gas exchange measurements, those that were conducted for bulb respiration were destructive. Therefore, calculating cumulative CO_2_ respired throughout the season could only be estimated with mean values, precluding us from estimating variation and, therefore, testing treatment effects statistically.

**Fig. 4:**
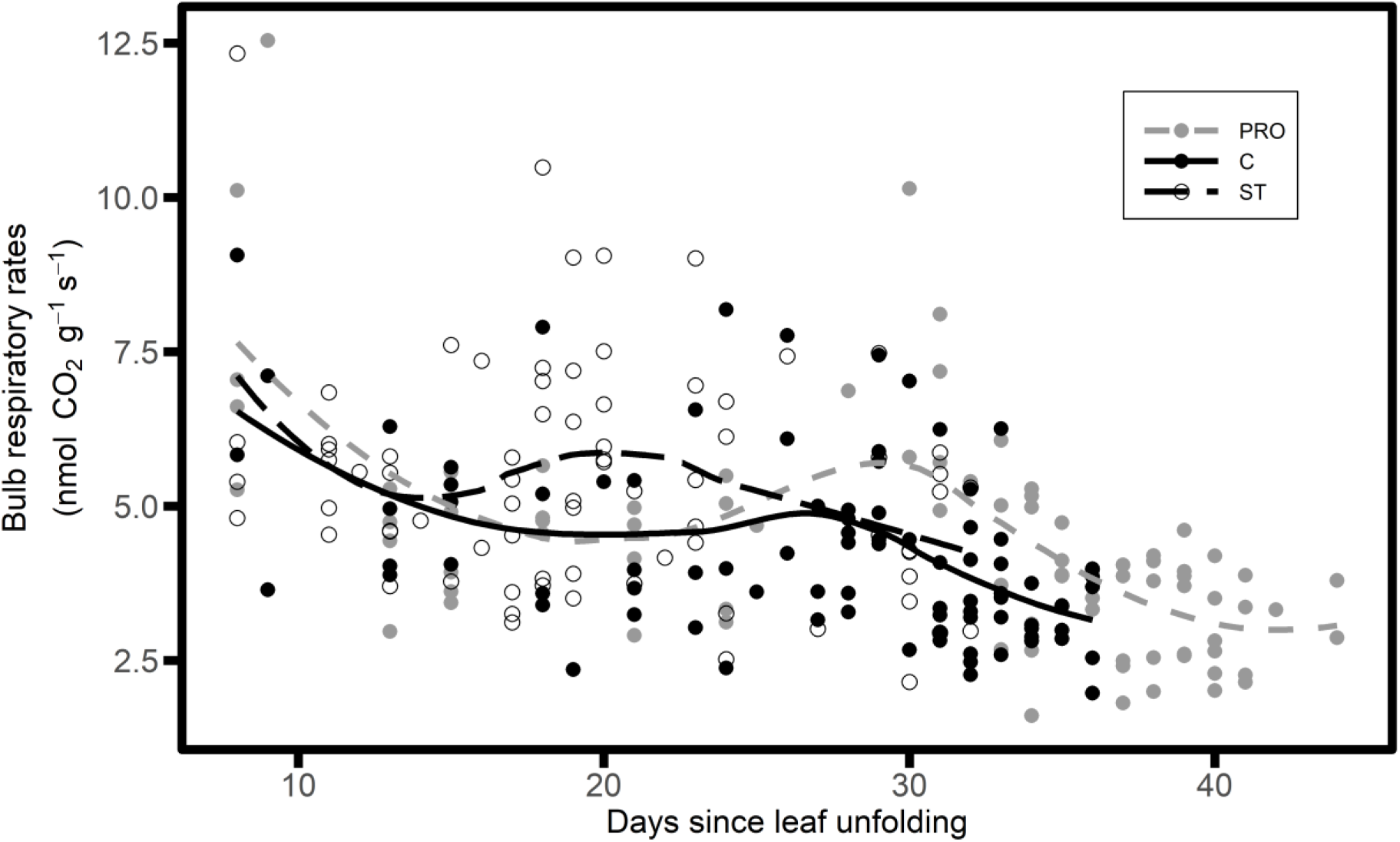
Bulb respiratory rates per bulb dry mass of *E. americanum* through time in plants subjected to foliar application of Promalin (PRO, grey), silver thiosulphate solution (STS, open circles) or water (Control, C, black). Each data point represents an individual plant and the lines follow a LOESS regression for each treatment.

**Table 2:** Results of repeated measures ANOVA testing the effects of treatments and time (days) on leaf chlorophyll fluorescence variables, gas exchange and C assimilation. *F*- values are presented along with statistical significance. *P*-values in bold are significant at *P* ≤ 0.05.

| Variables | Treatment | | Time | | Treatment × Time | |
| --- | --- | --- | --- | --- | --- | --- |
|  | *F* | *P* | *F* | *P* | *F* | *P* |
| ***Leaf*<sup>a</sup>** |  |  |  |  |  |  |
| Fv/Fm | 0.48 | 0.63 | 15.1 | **<0.001** | 2.41 | 0.093 |
| Fv'/Fm' | 0.15 | 0.86 | 74.3 | **<0.001** | 3.96 | **0.021** |
| ΦPSII | 0.73 | 0.50 | 140 | **<0.001** | 5.68 | **0.004** |
| qN | 0.48 | 0.63 | 85.4 | **<0.001** | 2.13 | 0.122 |
| qL | 0.98 | 0.40 | 91.5 | **<0.001** | 3.84 | **0.023** |
| Photosynthetic rates | 0.54 | 0.59 | 79.4 | **<0.001** | 3.14 | **0.046** |
| Respiratory rates | 1.19 | 0.33 | 1.58 | 0.211 | 2.09 | 0.127 |
| Total assimilated C per leaf area | 15.1 | **<0.001** | - | - | - | - |
| ***Bulb*<sup>b</sup>** |  |  |  |  |  |  |
| Respiratory rates | 0.55 | 0.58 | 3.73 | **0.003** | 0.78 | 0.671 |

<sup>a</sup>Degrees of freedom for Treatment: 2, Time: 1, interaction: 1; degree of freedom for the error term for Treatment: 13, Time: 183, interaction: 183.
<sup>b</sup>Degrees of freedom for Treatment: 2, Time: 6, interaction: 12; error term: 64.

Photosynthetic rates increased for the first 7 to 10 days following leaf unfolding and attained a maximum of 10.1 ± 0.5, 10.7 ± 0.3 and 11.3 ± 0.1 µmol CO_2_ m^-2^ s^-1^ for STS, Control and PRO plants, respectively (Fig. 5A). All three groups exhibited similar increase early in the season, given that treatments were applied when the peak in photosynthetic rates was attained. Photosynthetic rates then decreased until complete senescence, but at a steeper rate for STS compared to Control and PRO plants (Table 2; Fig. 5A). Given that leaf photosynthetic rates were measured on the same plants throughout the growing season, we could estimate total carbon assimilated by each plant. The three treatments differed in total C assimilated per leaf area, with values of 8.3 ± 0.7 mg C cm^-2^, 11.7 ± 0.7 mg C cm^-2^ and 14.3 ± 0.8 mg C cm^-2^ for STS, Control and PRO plants, respectively (Table 2).

**Fig. 5:**
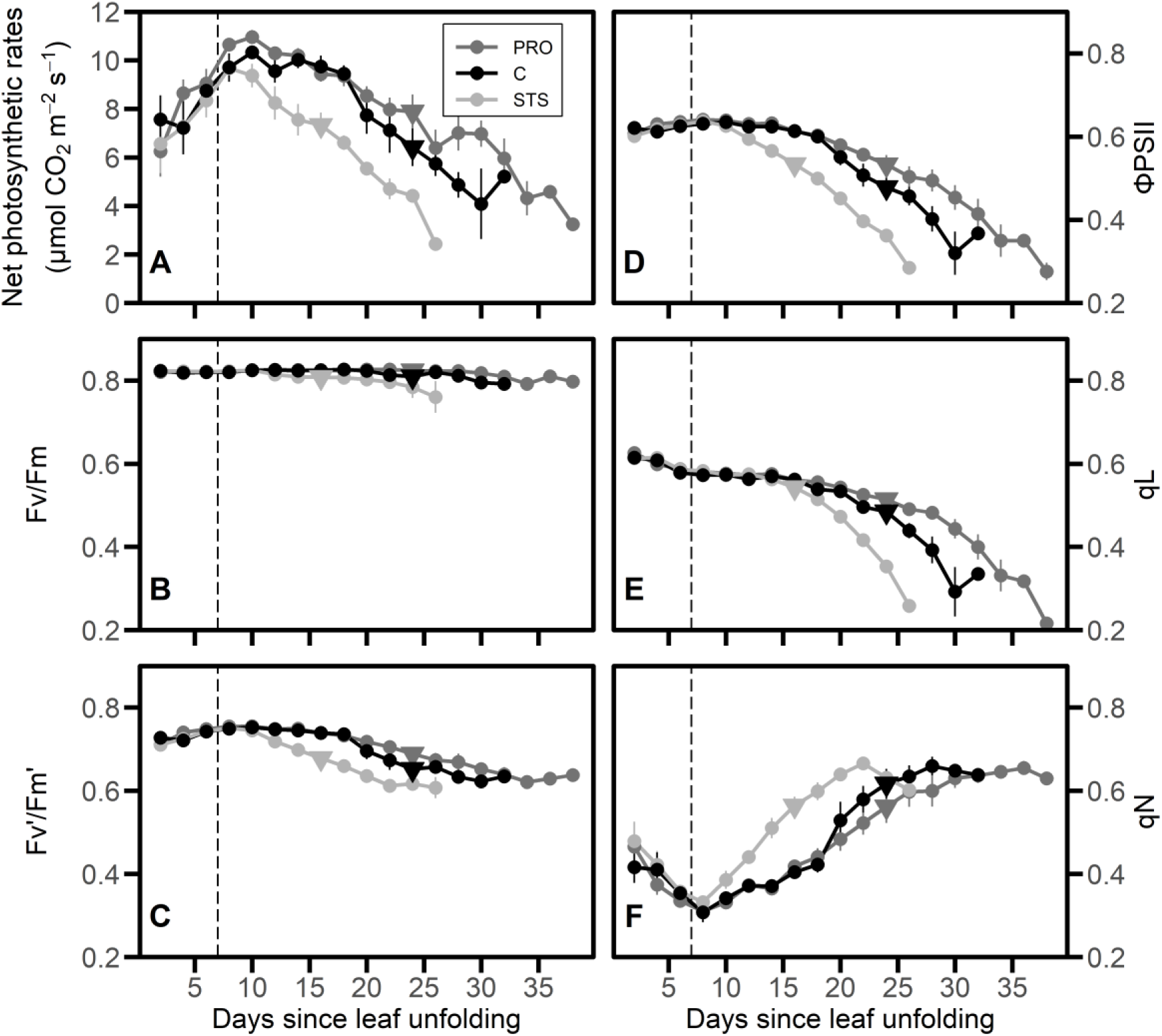
Evolution of photosynthetic rates (A), Fv/Fm (B), Fv’/Fm’ (C), ΦPSII (D), photochemical quenching (E), and non-photochemical quenching (F) through time in *E. americanum* plants subjected to foliar application of Promalin (PRO, grey), silver thiosulphate solution (STS, light grey) or water (Control, C, black). The vertical dashed line represents the time at which the first spraying occurred (7 days following complete leaf unfolding). Data points covered with a triangle indicate the beginning of leaf senescence within the three treatment groups. Means ± standard errors are presented (n = 6).

### Chlorophyll Fluorescence

Maximum efficiency of PSII photochemistry that was measured after dark adaptation, i.e., the Fv/Fm, remained high in all three treatments for the entire growing season in the green sections of the leaf (Fig. 5B). Fv/Fm values exhibited a slight reduction at senescence and tended to decline faster for STS plants. PSII maximum efficiency at 300 µmol photons m^-2^ s^-1^, which is denoted as Fv’/Fm’, showed an initial increase until 10 days after leaf unfolding; thereafter, it began to drop. The decrease that was observed in Control plants was intermediate between those of PRO and STS plants, where STS exhibited a decrease earlier than PRO plants (Fig. 5C). The actual efficiency of PSII at 300 µmol m^-2^ s^-1^ (ΦPSII) also exhibited a faster drop in STS plants as compared to Control and PRO plants (Fig. 5D). The fraction of ‘open’ PSII (qL) showed the same pattern as ΦPSII (Table 2, Fig. 5E); both qL and ΦPSII exhibited a stronger decrease through time than did Fv’/Fm’. The nonphotochemical quenching (qN) initially decreased up to 10 days after leaf unfolding; thereafter, it exhibited a continued increase until the leaf reached 25% senescence, after which it plateaued (Fig. 5F). All three treatments exhibited similar changes in qN over time (Table 2).

### Non-Structural Carbohydrate Accumulation

In the 6 days that had elapsed between the time that leaf unfolding was completed and the time where maximum photosynthetic rates were attained, bulb starch concentrations increased from 185 ± 29 mg g^-1^ to 780 ± 26 mg g^-1^ (Fig. 6A). In the days following treatment application, starch concentration decreased slightly, then recovered to values attained earlier on. This temporary drop in starch concentrations might be explained by the decline in photosynthetic rates which occurred at the same time. Starch concentrations did not differ among treatments, except at 5% senescence, where STS plants exhibited lower values (Table 3; significant treatment × stage interaction). This response may be explained by the fact that for STS plants, senescence began during the drop in starch concentration. Throughout most of the growing season, starch concentrations stayed near maximum values and starch content of the bulb continued to increase as bulb size increased. Starch content per leaf area ranged from 12.1 ± 0.9 mg cm^-2^ at maximum photosynthetic rates to 20.3 ± 3.7, 32.6 ± 1.6 and 32.8 ± 4.4 at final harvest for STS, Control and PRO plants, respectively (Table 3; significant treatment × stage interaction). This suggests that at any given time, additional cell growth was rapidly followed by starch synthesis to maintain maximum concentration values. Reducing sugars decreased at a rapid rate in the bulb between complete leaf unfolding (156 ± 30 mg g^-1^) and maximum photosynthetic activity (4.9 ± 1.6 mg g^-1^). Their concentrations then remained low until final harvest (Fig. 6B). Reducing sugar concentrations were slightly but significantly more elevated in Control plants than in the two other treatments (Table 3). Sucrose concentrations remained stable in the bulb until treatments were applied (Fig. 6C). Thereafter, sucrose concentrations decreased until final harvest for all three treatments, but remained higher for PRO plants as compared to STS and Control plants. The most abundant soluble sugar was sucrose, except at the very beginning of the growth period when reducing sugars were more abundant.

**Fig. 6:**
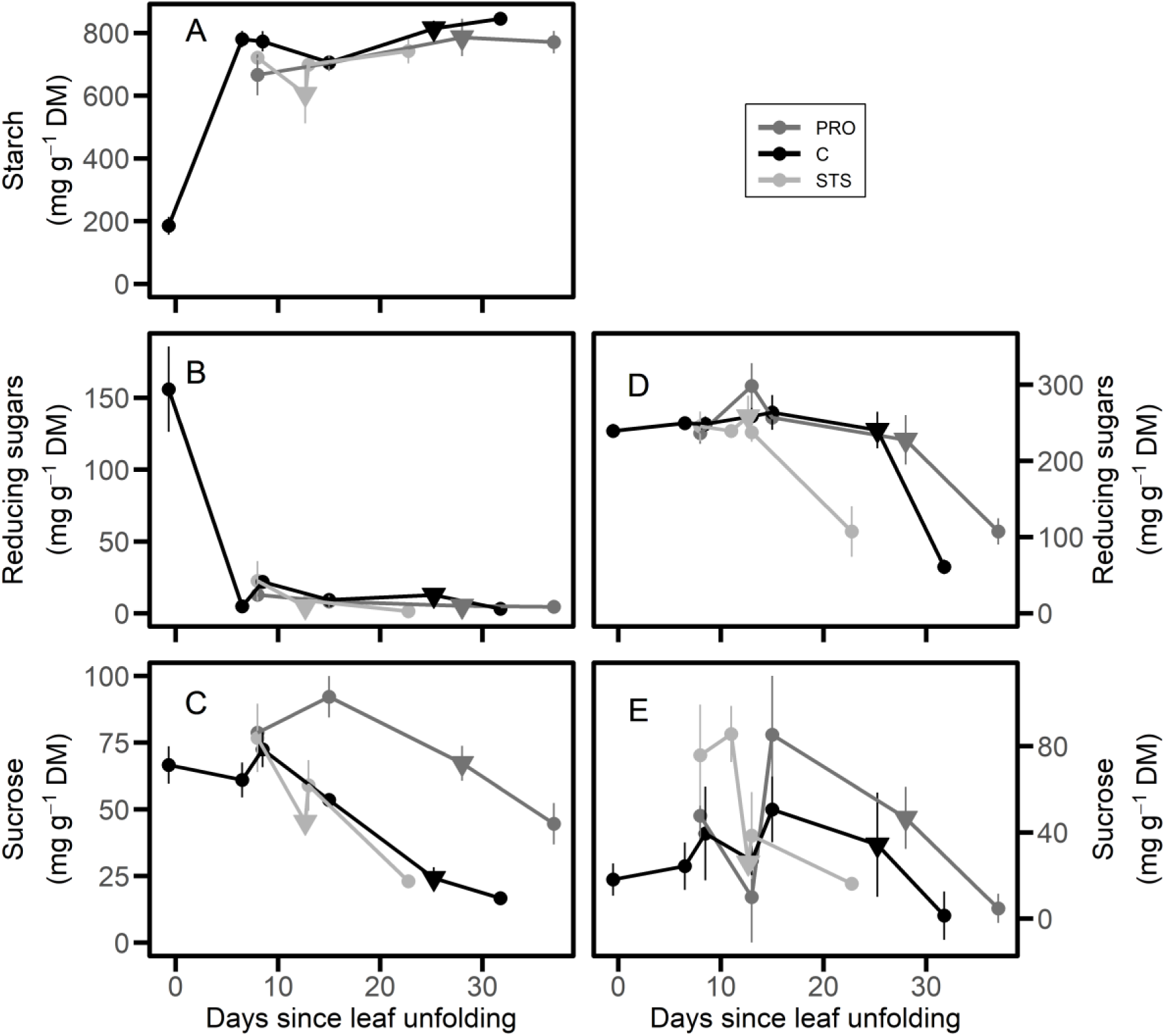
Starch (A), reducing sugar (B, D) and sucrose concentration (C, E) in bulbs (A, B, C) and leaves (D, E) of *E. americanum* receiving foliar applications of Promalin (PRO, grey) silver thiosulphate solution (STS, light grey), or water (Control, C, black). Treatments were applied 7 days after complete leaf unfolding (corresponding to the first data points for STS and PRO in panels A–E). Data points covered with a triangle indicate the beginning of leaf senescence within the three treatment groups. Means ± standard errors are presented (*n* = 4).

**Table 3:**
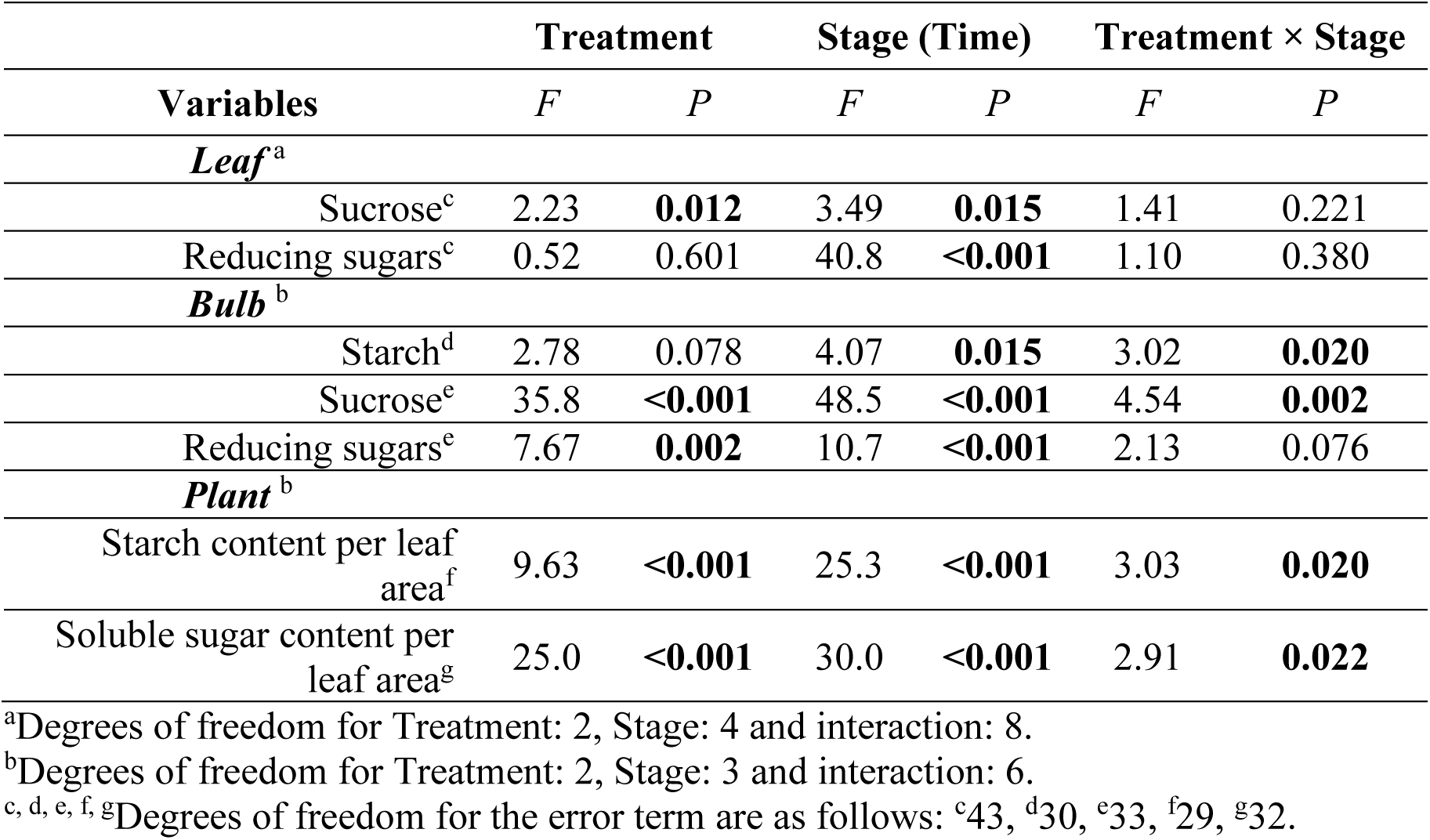
Results of two-way ANOVAs testing the effect of the treatments and developmental stages on non-structural carbohydrates in the leaves and bulbs. Total soluble sugar at the plant level (leaf and bulb) was also analyzed. *F*-values are presented along with statistical significance*. P*-values in bold are significant at *P* ≤ 0.05.

As reported previously by Gandin et al. (2009), we did not record any starch accumulation in the leaves. Similar amounts of reducing sugars were present in the leaves regardless of treatment (Fig. 6D; Table 3). Reducing sugar concentrations remained high throughout most of the season (250 ± 6 mg g^-1^), then dropped at final harvest as leaves reached complete senescence. HPLC analysis of sugar content in leaves of Control plants revealed that most of the reducing sugars were fructose and glucose (Supplementary Data 3). Furthermore, leaf glucose concentrations increased from the early developmental stages to the beginning of senescence (*R*^2^ = 0.57, *P* = 0.004, *df* = 9; Supplementary Data 3). Sucrose temporarily increased in concentration a few days ahead of the first sign of leaf senescence (15 days for Control and PRO, and 8 days for STS plants; Fig. 6E). Furthermore, sucrose concentrations were higher in PRO than in STS or Control plants. We estimated the total amount of soluble sugars per plant by summing leaf and bulb content and reporting it on a leaf area basis. Plant soluble sugar pools per leaf area were higher during most of the growing season in PRO plants, even at the end of leaf senescence (Fig. 7; Table 3). This is mainly due to the higher sucrose concentrations that were found in the bulb during most of the season (Fig. 6C).

**Fig. 7:**
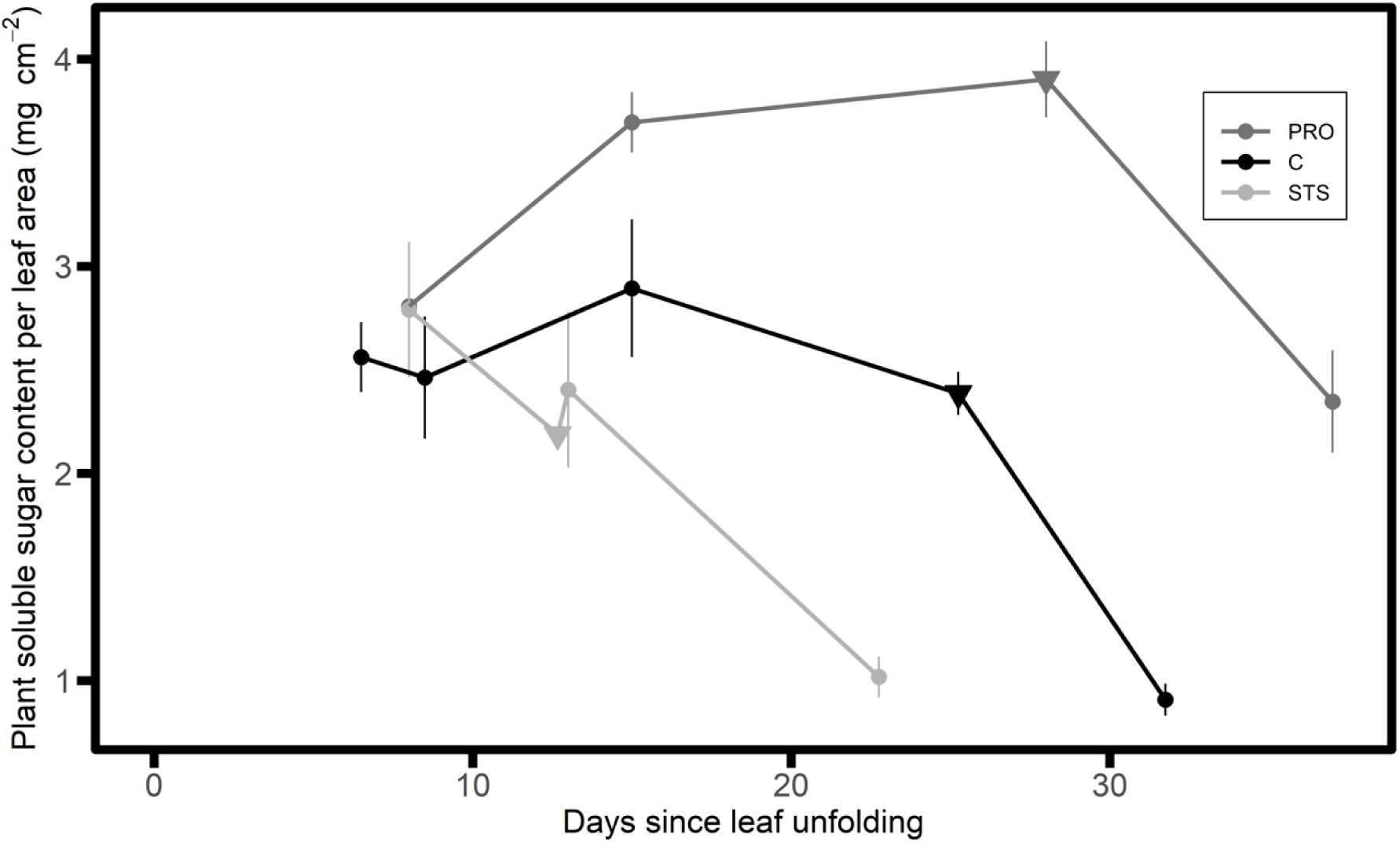
Total soluble sugar content in *E. americanum* plant (leaf and bulb) as expressed per leaf area for the three treatments: foliar application of Promalin (PRO, grey), silver thiosulphate solution (STS, light grey) or water (Control, C, black). Treatments were applied 7 days after complete leaf unfolding. Data points covered with a triangle indicate the beginning of leaf senescence within the three treatment groups. Means ± standard errors are presented (*n* = 4).

### Carbon Translocation

The first ^13^C pulse labelling took place when plants exhibited maximum photosynthetic rates (see Fig. 5A). Most ^13^C translocation occurred within the first 24 h, except in STS plants where bulb ^13^C concentrations continued to increase between 24 h and 48 h after pulse labelling (Fig. 8A, Table 4). At the second pulse labelling, which took place at the 5% leaf senescence stage, bulb ^13^C concentrations reached a maximum after 24 h in all three treatments, yet bulb ^13^C concentrations tended to continue to increase between 24 and 48 h in PRO plants (Table 4; Significant Labelling stage × Chase period × Treatment interaction) (Fig. 8B). Although it provided useful information regarding the translocation of newly assimilated C from the leaf to the bulb, the excess ^13^C per bulb dry mass does not account for bulb dilution due to the increase in size between first and second pulse labelling. This explains the much lower values that were observed at the second pulse labelling.

**Fig. 8:**
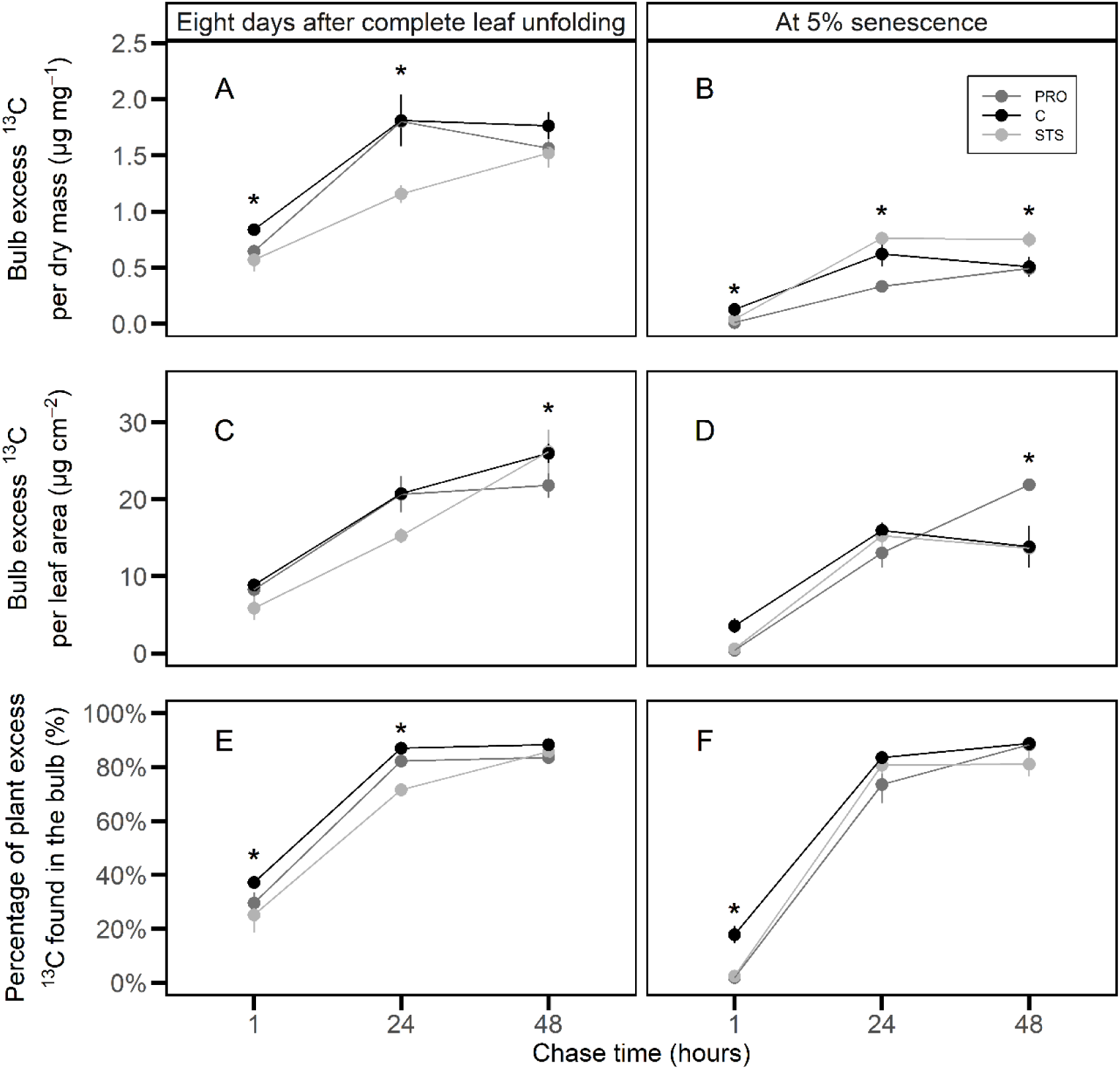
Bulb excess ^13^C per bulb dry mass (A, B) or per leaf area (C, D) and the fraction of plant excess ^13^C found in the bulb (E, F) at the end of the pulse-labelling period (1 h), 24 h and 48 h after pulse labelling. The first pulse labelling took place following the first foliar application (A, C, E) and the second pulse labelling at the beginning of leaf senescence (B, D, F) for plants treated with Promalin (PRO, grey), silver thiosulphate (STS, light grey) or water (Control, C, black). Asterisks indicate significant differences among treatments within a single chase period and labelling stage. Means and standard errors are presented (*n* = 4).

**Table 4:**
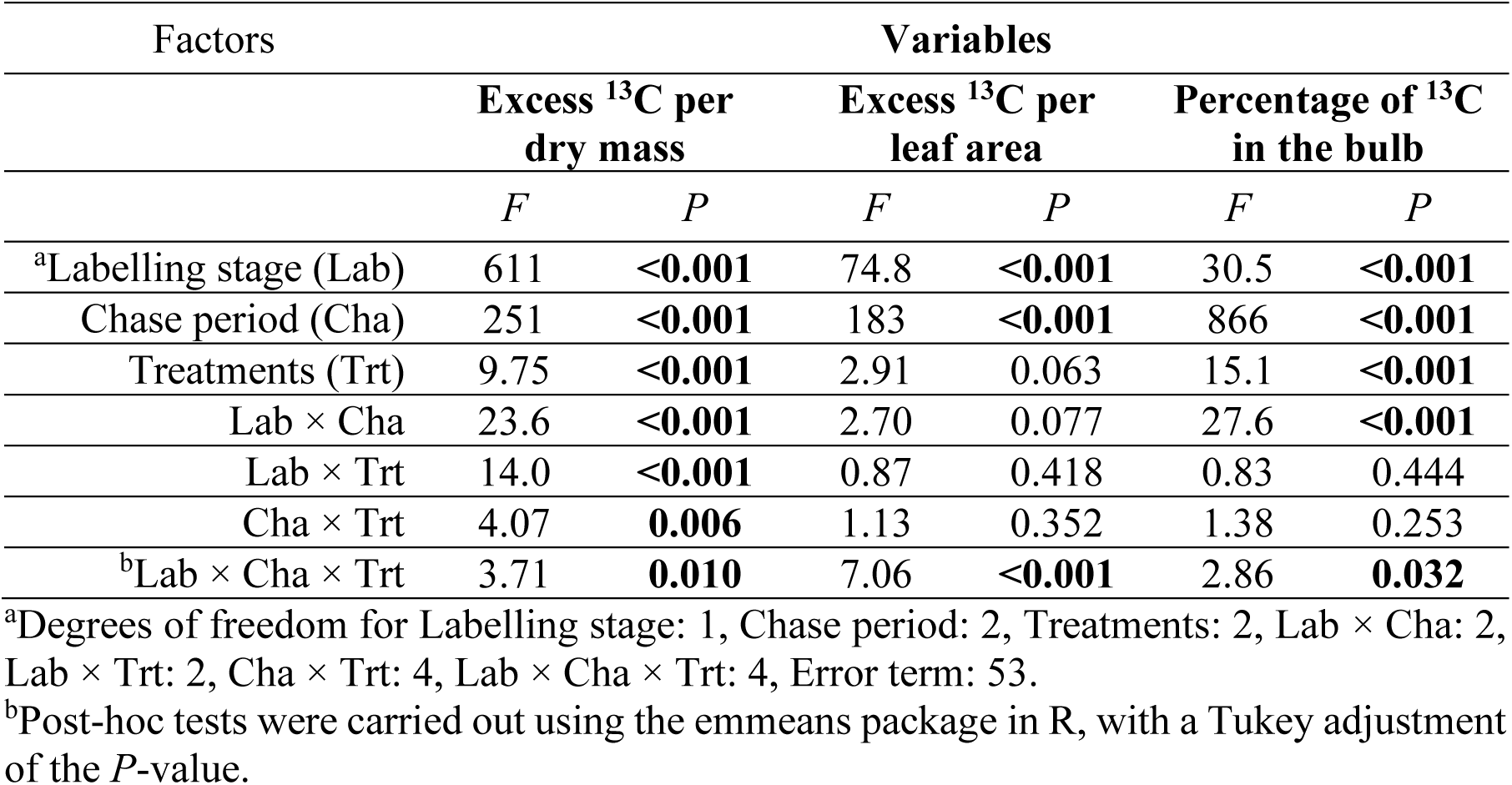
Results of three-way ANOVAs testing the effects of labelling stage (8 days after leaf unfolding vs. 5% leaf senescence), chase period and treatments on bulb excess ^13^C per dry mass or leaf area and the percentage of plant excess ^13^C found in the bulb. *F*-values are presented along with statistical significance. *P*-values in bold are significant at *P* ≤ 0.05.

| Factors | Variables |  |  |  |  |  |
| --- | --- | --- | --- | --- | --- | --- |
| | Excess $^{13}\text{C}$ per dry mass | | Excess $^{13}\text{C}$ per leaf area | | Percentage of $^{13}\text{C}$ in the bulb | |
| | $F$ | $P$ | $F$ | $P$ | $F$ | $P$ |
| <sup>a</sup>Labelling stage (Lab) | 611 | **<0.001** | 74.8 | **<0.001** | 30.5 | **<0.001** |
| Chase period (Cha) | 251 | **<0.001** | 183 | **<0.001** | 866 | **<0.001** |
| Treatments (Trt) | 9.75 | **<0.001** | 2.91 | 0.063 | 15.1 | **<0.001** |
| Lab × Cha | 23.6 | **<0.001** | 2.70 | 0.077 | 27.6 | **<0.001** |
| Lab × Trt | 14.0 | **<0.001** | 0.87 | 0.418 | 0.83 | 0.444 |
| Cha × Trt | 4.07 | **0.006** | 1.13 | 0.352 | 1.38 | 0.253 |
| <sup>b</sup>Lab × Cha × Trt | 3.71 | **0.010** | 7.06 | **<0.001** | 2.86 | **0.032** |

<sup>a</sup>Degrees of freedom for Labelling stage: 1, Chase period: 2, Treatments: 2, Lab × Cha: 2, Lab × Trt: 2, Cha × Trt: 4, Lab × Cha × Trt: 4, Error term: 53.
<sup>b</sup>Post-hoc tests were carried out using the emmeans package in R, with a Tukey adjustment of the $P$ -value.

^13^CO_2_ assimilated during pulse labelling is also influenced by plant leaf area. To overcome these issues, we also expressed bulb ^13^C content per leaf area. During the first pulse labelling, the amount of ^13^C that was translocated to the bulb per leaf area continued to increase between 24 h and 48 h in Control and STS plants, whereas in PRO plants, it reached a maximum after 24 h (Fig. 8C). At the second pulse labelling, the amount of ^13^C that was translocated to the bulb per leaf area reached its maximum at 24 h in Control and STS plants, whereas it continued to increase in PRO plants between 24 and 48 h (Fig. 8D).

The same percentage of assimilated ^13^C was found in the bulb 48 h after pulse labelling in all three treatments and during both pulse labellings. This implies that within two days, around 85% of what had been assimilated in the leaves was now located in the bulbs, irrespective of treatment or leaf age. The increasing percentage of plant excess ^13^C found in the bulb that was observed between 24 h and 48 h in STS plants during the first pulse labelling and in PRO plants during the second pulse labelling, thus suggests a slower translocation rate towards the bulb (Fig. 8E, 8F). At the first pulse-labelling, the percentage of ^13^C in the plant that was located in the bulb after 24h was lower for STS (72.6 ± 2.3%) than for the Control plants (87.0 ± 1.8%). In terms of sucrose, this difference represented 1.75 ± 0.42 mg that was not yet transferred in the bulb. In the leaves, this delay in translocation would translate into an increase of 46 ± 12 mg g^-1^ during the days following the first pulse labelling, and would be consistent with leaf sucrose concentrations that were reported in Fig. 6E. The sucrose that had not yet been translocated to the bulb 24 h after the end of the second pulse labelling in PRO plants amounted to 0.91 ± 0.65 mg. This would correspond to an increase of 28 ± 21 mg g^-1^ in the leaves, which is similar to the difference that was shown previously (Fig. 6E).

Given that 85% of all C that was assimilated by the plant was eventually translocated to the bulb, irrespective of leaf age (Fig. 8E, 8F), we can estimate the actual amount of daily C that was translocated to the bulb per leaf area (mg C cm^-2^ day^-1^) by converting photosynthetic rates (Fig. 5A) to daily C fixed per leaf area, and then multiplying this value by 0.85 (Fig. 9). By converting bulb respiratory rates (Fig. 4) into daily C respired per leaf area (mg C cm^-2^ day^-1^), we can then subtract this value from the amount of C that was translocated to the bulb and estimate the amount of C that is available for growth. The amount of C that was translocated followed the decrease in photosynthetic rates (Fig. 5A), which is expected since 1) leaf area did not change once leaf unfolding was completed, and 2) the percentage of C that was translocated remained constant (Fig. 8C, 8F). Bulb respiratory rates per leaf area increased early in the season, then reached a temporary plateau 16-19 days following complete leaf unfolding for both Control and PRO plants (Fig. 9B, 9C). A burst of bulb respiratory rates was observed at 28-31 days in these two treatment groups and, when reported on a daily basis, accounted for most of the daily C translocated to the bulb. This leads to nearly no carbon being available for growth near the end of the growing season. In STS plants, bulb respiratory rates exhibited a continuous increase as the bulb increased in size, to the point that no carbon remained available for growth 20 days following leaf unfolding. These results agree with the growth curves (Fig. 3) that were reported for the three groups of plants. The increase in bulb respiratory rates may be the result of increased C availability towards the end of the season when bulb growth had ceased. However, our results do not indicate the cause of growth cessation, but only that we can account for all C that has been translocated throughout the season.

**Fig. 9:**
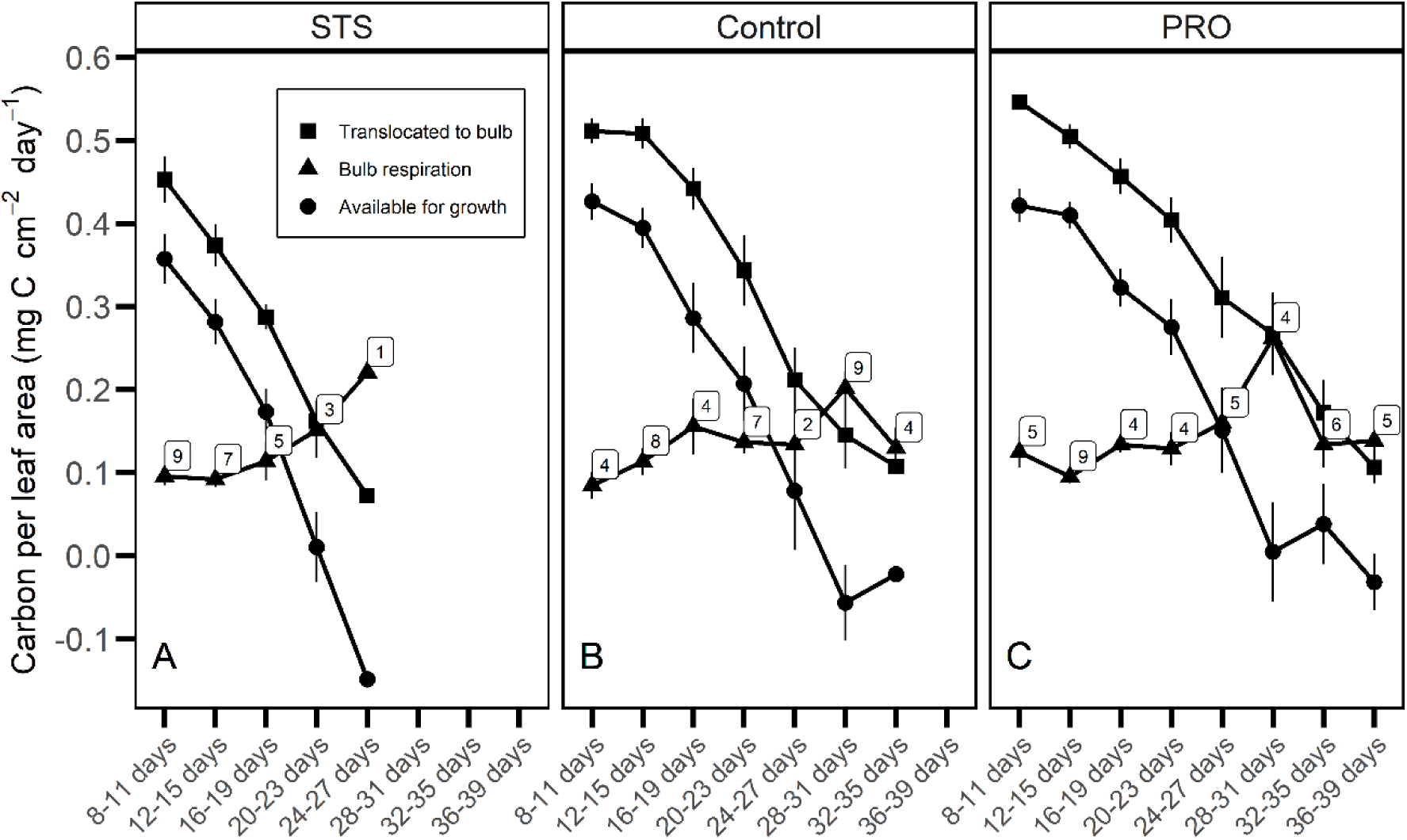
Daily carbon per leaf area that was translocated to the bulb (square), used for bulb respiration (triangle) or available for growth (circle) for *E. americanum* plants treated with Promalin (PRO), silver thiosulphate (STS), or water (Control, C). Each data point represents the mean (± standard error) of the daily carbon per leaf area that is translocated, respired or available for growth during a four-day interval. The number of plants that were measured per treatment during this interval is indicated for the bulb respiration data (*n* = 6 for translocation).

## Discussion

### Leaf Longevity, Photosynthetic Activity and Growth

In spring geophytes, warm temperatures decrease both bulb growth and leaf longevity (Badri *et al*., 2007; Bernatchez and Lapointe, 2012). Yet, growth decline at warmer temperatures can only partially be explained by the shorter leaf life span, at least in *E. americanum* (Gandin *et al*., 2011b). The increased leaf life span that was observed in Promalin-treated plants allowed more C assimilation over the season. Nevertheless, enhancing leaf life span had no significant effect on bulb growth at warm temperatures. We can conclude that the plant was incapable of converting this extra assimilated C into more biomass. Reducing leaf life span, however, did affect plant growth. Plants treated with STS exhibited a sharper decline in their photosynthetic rates than did the Control plants. This reduction, when combined with their shorter leaf life span, reduced the total amount of C that was assimilated during the season and restricted bulb growth of STS compared to that of Control plants.

Photosynthetic rates decreased over time in all three treatment groups as the increase in bulb mass slowed. Given that both occurred concurrently, we cannot distinguish which one is causing the other to change (Adams *et al*., 2013). Yet, fluorescence data support the hypothesis of the build up of a sink limitation over time. In both Control and treated plants, PSII operating efficiency (ΦPSII) exhibited a continuous decrease over time, concurrent with decreasing photosynthetic rates. This response occurred with very little change in Fv/Fm, indicating that the photosynthetic apparatus remained intact. Furthermore, the continuous decrease that was observed for ΦPSII and photochemical quenching (qL) is accompanied by a constant increase in non-photochemical quenching (qN), strongly suggesting that photosynthetic activity is down-regulated and that the photosystems must direct some of the energy towards heat dissipation (Pammenter *et al*., 1993). Therefore, a feedback inhibition of the photosynthetic rates due to decreasing sink demand is supported by the fluorescence data, as has been reported previously not only in *E. americanum* (Badri *et al*., 2007; Gandin *et al*., 2011b), but also in young apple (*Malus × domestica* Borkh.) trees (Wunsche *et al*., 2005).

It has been shown that the alternative respiratory pathway may be induced to maintain a balance between photosynthesis and respiration in leaves of tobacco (*Nicotiana tabacum* L.) plants that are grown under elevated CO_2_ (Dahal and Vanlerberghe, 2018). In a previous study on *E. americanum* where source strength was modulated daily by exposing the plant to high CO_2_ concentrations (Gandin *et al*., 2009), the bulb used the alternate respiratory pathway to cope with the imbalance between the amount of C coming from the source and its capacity to use it. Additional assimilated C during the day was thus respired, given that it could not be used for additional bulb growth (Gandin *et al*., 2009). Nevertheless, by extending the growing season rather than the daily amount of C that is translocated to the bulb, we expected the bulb to continue growing and to achieve a larger size, as reported in *A. tricoccum*, another spring ephemeral (Dion *et al*., 2017). Yet, due to the steady decline in photosynthetic rates and the concurrent increase in bulb respiratory rates through time, increasing leaf longevity of *E. americanum* did not translate into more C accumulation in the bulb. Most of the C that was translocated to the bulb was accounted for by maintenance respiration towards the end of the season. This resulted in comparable bulb sizes for Control and PRO plants. The burst of bulb respiratory rates that was observed near the end of the growing season (Fig. 9) might represent a mechanism for coping with daily source-sink imbalance or sink inability to grow larger (Gandin *et al*., 2009; Campany *et al*., 2017).

A recent publication by Greco et al. (2019) revealed a negative relation between *E. americanum* abundance and late canopy closure. This surprising observation is contrary to those of various other spring ephemerals, where growth is usually enhanced under a late canopy closure or reduced under early tree leaf-out (Kim *et al*., 2015; Dion *et al*., 2017; Heinrichs *et al*., 2018). Abundance might be controlled, however, by factors differing from those affecting plant growth, e.g., factors that favour vegetative propagation.

### Developmental Phases and Growth

Why do bulbs stop growing before leaf senescence occurs? Final organ size is a function of its cell number and individual cell size. Sink initiation and expansion are mainly associated with invertase activity, whereas maturation and storage are driven by sucrose synthase (Koch, 2004). The major storage constituent of *E. americanum* bulb is starch, which makes up nearly 80% of bulb dry mass. Starch accumulation, which normally should occur during the maturation phase, starts much earlier in the growing season in *E. americanum*, and earlier at higher than at lower temperatures (Gandin *et al*., 2011b). Sucrose synthase activity is indeed present early in the season, while invertases (cell wall and vacuolar) are still active (Gandin *et al*., 2011b). In concordance with these results, cell elongation continues during starch accumulation, which means that both expansion and maturation are concomitant processes in bulbs of spring ephemerals (Gandin *et al*., 2011b) and most likely in corms of spring geophytes as well (Badri *et al*., 2007). We do not know, however, when cell division stops. Bulb cells reached a smaller size at the end of the season at a temperature of 18 °C compared to 12 °C and 8 °C (Gandin *et al*., 2011b), but the difference in cell size did not fully explain the reduction in bulb mass at 18 °C compared to the two colder temperatures. Furthermore, assuming a nearly identical bulb density, as suggested by their starch and soluble sugar concentrations, the number of cells that are contained in the bulb at 18 °C would be roughly 35% lower than that at the two cooler growth temperatures. There is usually little overlap between cell division and elongation in leaves (Gonzalez *et al*., 2012) or roots (Beemster *et al*., 2003). Yet, in potato tubers, cell division continues throughout tuber growth (Chen and Setter, 2012). If the same is true for bulbs, this would mean that all three activities – cell division, cell elongation and carbohydrate accumulation – occur concomitantly. It appears that the effect of higher temperature on *E. americanum* bulb cell size, and potentially on cell number as well, cannot be explained by the shorter growing season, since PRO plants produced bulb of similar size as Control plants. Cell proliferation and enlargement appears to be reduced at warmer temperatures by a mechanism other than carbon availability.

Interestingly, we found that bulb growth is more strongly associated with leaf area than with the previous autumnal bulb mass. This is surprising, given that source strength has been demonstrated to have no influence on bulb growth in this species (Gandin *et al*., 2009). The strong correlation between bulb size and leaf area could mean that maximal bulb size is set conjointly with leaf development. In spring geophytes such as *E. americanum*, leaf cell division within the new bud occurs the previous summer, whereas cell elongation takes place slowly during the autumn and winter, then rapidly in spring after snowmelt (Grime and Mowforth, 1982). Leaf mesophyll of spring ephemerals is already partially differentiated while the leaf tissues are still underground in late winter and contains proplastids, which will develop into fully functional chloroplasts a few days after being exposed to light (Mamushina *et al*., 2002). At that point, the leaf is connected to the mother bulb, which provides carbohydrates for its rapid growth. The new bulb (daughter bulb) becomes apparent after the leaf has completed its expansion (Dong, 2020). If this strong correlation between leaf area and new bulb mass indeed indicates the presence of a common set of regulatory factors, these factors would thus have to play a role during the previous summer at the time when the leaf is being initiated.

It has been posited that developmental processes prior to starch accumulation in wheat grains limit grain size as shown by Fahy *et al*. (2018) for 21 *Triticum aestivum* genotypes. The same could be taking place in *E. americanum* bulbs. Increasing the length of the maturation period does not influence growth if earlier developmental periods are not conjointly extended. This idea is supported by a study on *E. americanum* in which a shorter photoperiod led to late starch saturation of the sink and delayed leaf senescence, without altering final bulb biomass, compared to a longer photoperiod (Gandin *et al*., 2011a). Bulb division and expansion phases that take place at the beginning of the growing season might dictate the final size that the bulb can attain, even though both activities appear to continue throughout the season. Therefore, the bulb could not get bigger despite delaying leaf senescence, but it might stay smaller if the senescence signal occurs earlier, as shown by plants treated with STS.

### Carbohydrate Signalling

Leaf glucose sensing by hexokinase leads to a decrease in photosynthetic rates and accelerated senescence (Quirino *et al*., 2000). In the current study, no increase in reducing sugar content occurred in the leaves at the time senescence became apparent (Fig. 6D). Nevertheless, glucose concentrations may have increased while other reducing sugars decreased in concentration to the extent that total reducing sugar concentrations in the leaves remained relatively constant through time. Indeed, leaf glucose concentrations of Control plants (Supplementary Data 3) increased from the early stages to the beginning of leaf yellowing, suggesting a potential role of glucose in the induction of leaf senescence or in the decline in photosynthetic rates.

Sucrose concentrations exhibited more fluctuation than that of reducing sugars throughout the season, and we can clearly see a peak in sucrose in the leaves of STS and PRO plants (Fig. 6E) at the time leaf photosynthetic rate was at its maximum values (Fig. 5A). Sucrose cleaving enzymes, i.e., sucrose synthase or invertase, play a signalling role in plant development by modulating the availability of hexoses (Koch, 2004). Sucrose can also act as a signal on its own without acting on hexose concentrations and signalling (Tognetti *et al*., 2013). High concentrations of sucrose induce an increase in trehalose-6- phosphate (Tre6P) concentrations, which in turn acts as a signalling molecule that induces a reduction in photosynthetic rates (Lunn *et al*., 2014). Although bulb growth rate was maximum at that time (Fig. 3), it appears that the photosynthetic rates were higher than the sink capacity, which led to a temporary high level of sucrose until leaf photosynthetic rates had adjusted. Thereafter, it appears that the photosynthetic rates were regularly down- regulated with no transient increase in either reducing sugars or sucrose concentrations. Most likely, the first down-regulation induced a signalling pathway that remained active and which ensured fine-tuning of the source-sink relationship. A metabolomic study in *E. americanum* has reported an increase in glucose-6-phosphate and fructose-6-phosphate in the leaves at the time they attained maximum photosynthesis (Dong, 2020). These sugars might also play a role in initiating this signalling pathway.

Reducing sugars were only abundant early on in the bulb. Thereafter, sucrose was the main soluble sugar and its concentration kept decreasing, which was likely due to the dilution effect caused by the large accumulation of starch. These results agree with those of Gandin et al. (2011b) and suggest 1) that sucrose is the main sugar being translocated from leaf to bulb and 2) that the conversion from sucrose to starch occurs rapidly, thereby avoiding transient accumulation of glucose or fructose. Similar patterns of starch accumulation in the storage organ have been observed in the spring ephemeral mayapple (*Podophyllum peltatum* L.) (Constable *et al*., 2007). In tubers of potato (*Solanum tuberosum* L.), the intermediates of sucrose-starch conversion are also relatively stable during starch filling, due to the reversibility of sucrose breakdown by sucrose synthase (Geigenberger, 2003). Cells achieved maximum starch concentration very early, which also supports the hypothesis that starch synthesis occurs at a very fast rate. Activity of AGPase, one of the enzymes involved in starch synthesis, remains high throughout the season (Gandin *et al*., 2011b), also supporting the hypothesis of a strong starch synthesis capacity. A recent metabolomics study on *E. americanum* reported a transient increase in glucose-6-phosphate in the bulb at the same time that this metabolite increased in the leaf, i.e., at maximum photosynthesis (Dong, 2020). The ‘smooth’ regulation of photosynthetic rates following this initial imbalance, and the large accumulation of starch, most likely masked any potential transient increase in sucrose. Indeed, Gandin et al. (2011b) and Dong et al. (2018) reported soluble sugars as a function of bulb dry mass from which starch content had been subtracted in an attempt to estimate sugar concentrations in the cytosol. Both studies indicated a clear transient peak in soluble sugars a few days prior to the first sign of leaf senescence at the warmer temperature (i.e., 18 °C). Soluble sugars temporarily accumulate early in the season in the leaf as photosynthetic rates reached their maximum values, inducing a reduction in photosynthetic rates. Later on, they likely accumulate again in the cytosol of bulb cells, but this time, the feedback mechanism induces leaf senescence.

### Promalin and Silver Thiosulphate Modulation of Source-Sink Signalling

Ethylene production in mature leaves is associated with the onset of leaf senescence (Koyama, 2014). Yet, the ethylene synthesis inhibitor (AOA) and action inhibitor (STS) that were tested prior to this study stimulated leaf senescence in *E. americanum* instead of delaying it (Supplementary data Fig. S1). Ethylene can induce higher photosynthetic rates and greater plant growth in tobacco and mustard (*Brassica juncea* L. Czern & Coss.) by decreasing sensitivity to glucose (Tholen *et al*., 2007; Iqbal *et al*., 2011). The inhibition of ethylene action by STS could have induced the observed drop in *E. americanum* photosynthesis by not inactivating the hexokinase signal. Yet, the hexokinase response does not account for the sucrose increase in the leaf. Differences in the percentage of total ^13^C that were found in the bulb between Control and STS plants 24 h after the first pulse labelling would translate into a build-up of about 46 mg g^-1^ of sucrose in the leaves of STS plants. Results of the sugar assay corroborate the increased level of sugar in the leaf at that specific moment. We posit that one of the effects of STS was to inhibit sucrose translocation from the leaf to the sink shortly after foliar application, leading to a momentary high sucrose concentration in the leaf. This sucrose peak in the leaf, together with sustained sensitivity to glucose, likely caused the earlier senescence and stronger photosynthetic rate declines in this treatment, possibly through the Tre6P signalling pathway (Lunn *et al*., 2014). Whether sugar translocation may be slowed by the inactivation of the ethylene response or by other effects of STS remains unknown, given that there are few records regarding the effect of STS on sugar metabolism in the literature, and these mainly focus on its role in prolonging the life of cut flowers (Meir *et al*., 2013; Razali *et al*., 2013). Ethylene has also been shown to induce sucrose transporter gene expression in grape berries (*Vitis vinifera* L.) and rubber tree (*Hevea brasiliensis* M<u>ü</u>ll.Arg.) (Chervin *et al*., 2006; Dusotoit-Coucaud *et al*., 2009). The application of STS may have inhibited the synthesis of some sucrose transporters, by inactivating the response of the plant to ethylene. The lack of transporters when source strength exceeds sugar transport capacities may have led to the momentary increase in sucrose concentrations in the leaves, which was sufficient to induce leaf senescence.

The effect of reduced leaf duration on final bulb mass that was caused by STS is not consistent with previous studies where source activity was either reduced by continuous exposure to ozone (Gandin *et al*., 2009), or by low PPFD (Gandin *et al*., 2011a). In both of these studies, the reduction in source activity did not affect final bulb mass. The early induction of leaf senescence that was caused by STS might play a more important role in the final bulb mass than the reduced C assimilation that resulted from this early senescence. Indeed, leaf senescence might induce changes in the bulb, which end bulb growth prematurely. For example, commercial production of potatoes involves the application of herbicides to induce shoot senescence along with tuber maturation (Bethke and Busse, 2010). Although the relationship between leaf senescence and tuber maturation is not fully understood, we can posit that inducing premature leaf senescence through STS has hastened the developmental process in *E. americanum* bulbs, leading to smaller final bulb sizes.

Senescence is delayed by cytokinins through the activation of cell-wall invertase (Lara *et al*., 2004). The activation of cell-wall invertase in a weak source, such as the leaf at the end of the growing season, leads to a futile cycle of sucrose transport (Zwack and Rashotte, 2013). Instead of entering the companion cell via a sucrose transporter, sucrose is hydrolyzed into glucose and fructose by the cell-wall invertases in the apoplast. The hexoses are then returned to the phloem parenchyma and re-synthesized into sucrose. This illusion of a high sink demand caused by the application of cytokinins on leaves, delays senescence. The cycle of sucrose re-synthesis reduced sucrose translocation towards the bulb, as observed during ^13^C pulse-labelling at 5% senescence. Slowing sucrose translocation increases sucrose concentrations in the leaf, in the same way as with STS during the first pulse labelling. The increased sucrose concentrations in leaves of PRO plants possibly act as a competing signal for cell-wall invertase through the sucrose/Tre6P signalling pathway, thereby leading to the reduction of photosynthetic rates despite the action of cytokinins.

## Conclusion

Under natural conditions, climate change is already causing an increase in spring temperatures, which shorten the leaf life span of numerous spring flowering herbs (Augspurger and Zaya, 2020). Our findings suggest that lengthening the maturation phase in *E. americanum* did not affect final bulb size under warm temperatures, even if cell size had the potential to keep increasing as shown with plants grown at lower temperatures. A strong feedback inhibition of photosynthesis and the upregulation of bulb respiration occurred concomitantly with a slowing of bulb growth at the end of the season for *E. americanum*. However, when senescence occurred prematurely, as was observed following exposure to STS, bulb growth was negatively affected. Therefore, a combination of warmer temperatures and earlier tree leafing-out would strongly reduce growth of spring ephemerals and eventually their abundance. One potential factor that might help their survival is early snowmelt, which could hasten their sprouting in early spring (Routhier and Lapointe, 2002), much more so than for trees which are influenced by a more complex set of factors (Raulier and Bernier, 2000; Ladwig *et al*., 2019). Nevertheless, warmer spring would affect the capacity of the plant to accumulate carbohydrates in its perennial organ, limiting its growth regardless of the duration of the high light period between its leaf unfolding and tree leaf emergence and development.

## Supplementary Data

Fig. S1: Leaf life duration of *E. americanum* sprayed with silver thiosulfate (STS), aminooxyacetic acid (AOA), benzylaminopurine (BAP), Promalin (PRO) or water (C), during a preliminary trial. Different letters indicate differences among treatments following one-way ANOVA (*F*_4,58_ = 54.7; *P* < 0.001) and post-hoc Tukey tests.

Table S1: Results of Pearson’s product moment correlation (*r*) analyses between plant leaf area and time elapsed between complete leaf unfolding and either the first visual observation of leaf 5% or 100% yellowing for *E. americanum* plants that were treated with silver thiosulfate (STS), water (C) or Promalin (PRO).

Table S2: Results of linear regression between autumnal bulb mass, leaf area and bulb mass at final harvest for *E. americanum* plants that were treated with silver thiosulfate (STS), water (C) or Promalin (PRO).

Supplementary Data 3: Methods, data (Table S3) and linear regression between leaf glucose concentrations (analyzed by HPLC-RID) and days since leaf unfolding (Fig. S2).

## Acknowledgements

The authors thank Steeve Pepin for his advice on ^13^C labelling protocol, for lending us some of his equipment and for providing comments on a first draft of the manuscript, Guy Samson for providing comments on a first draft of the manuscript, Gaétan Daigle for statistical consultation, and William F.J. Parsons for English revision of the text.

## Authors Contributions

HB designed and conducted the different experiments, analysed data and wrote the manuscript; LL planned the study, contributed resources and facilities and wrote the manuscript.

## Data availability

The data supporting the findings of this study are available from the corresponding author (Line Lapointe) upon request.

