## Supplementary data for "Bulb growth potential is independent of leaf longevity for the spring ephemeral *Erythronium americanum* Ker-Gawl"

Département de biologie and Centre d'étude de la forêt, Université Laval, Québec, Québec, Canada. G1V0A6

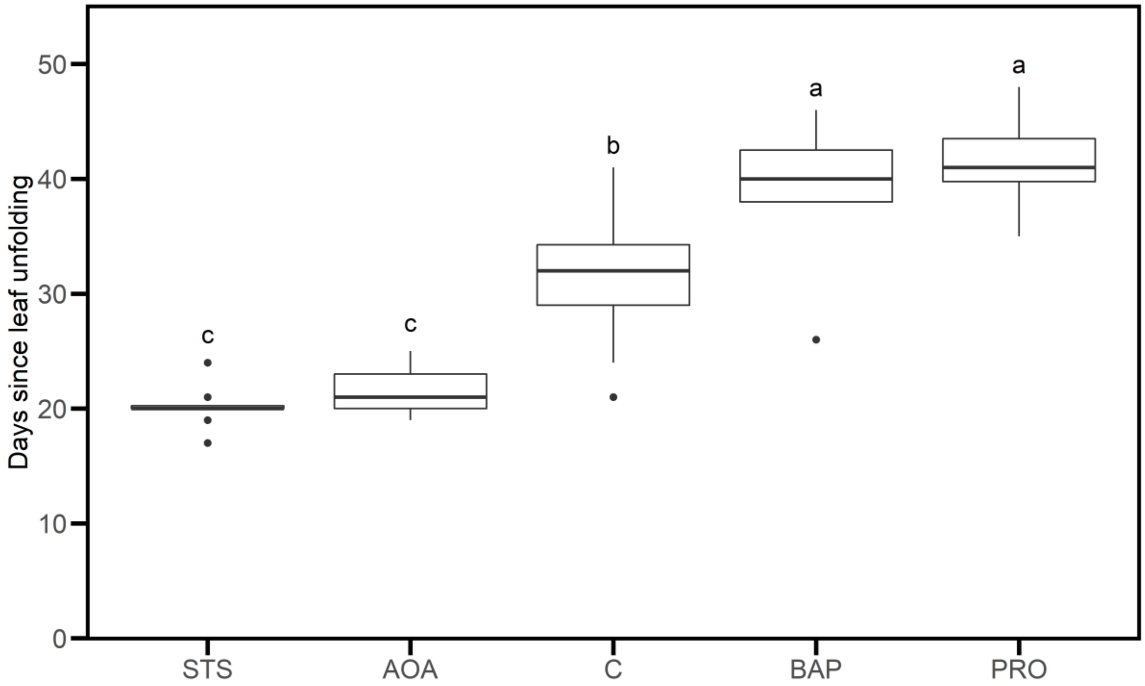


**Fig. S1**: Leaf life duration of *E. americanum* sprayed with silver thiosulfate (STS), aminooxyacetic acid (AOA), benzylaminopurine (BAP), Promalin (PRO) or water (C), during a preliminary trial. Boxes show interquartile range (IQR) with median indicated by horizontal line. Whiskers are ± 1.58 × IQR/√n. Outliers are shown as individual points. Different letters indicate differences among treatments following a one-way ANOVA ($F_{4,58} = 54.7$; $P < 0.001$) and post-hoc Tukey tests.

**Table S1**: Results of Pearson's product moment correlations between plant leaf area and time elapsed between complete leaf unfolding and either the first visual observation of leaf 5% or 100% yellowing for *E. americanum* plants treated with silver thiosulfate (STS), water (C) or Promalin (PRO).

| | **STS**[a] | | **C** | | **PRO** | |
|---|---|---|---|---|---|---|
| **Variables** | *r* | *P* | *r* | *P* | *r* | *P* |
| 5% senescence and leaf area | -0.15 | 0.33 | -0.12 | 0.38 | -0.19 | 0.25 |
| 100% senescence and leaf area | -0.07 | 0.66 | -0.01 | 0.95 | -0.10 | 0.55 |

[a]Degrees of freedom of the error term for STS: 42, C: 50, PRO: 37.

**Table S2**: Results of linear regression between autumnal bulb mass, leaf area and bulb mass at final harvest for *E. americanum* plants treated with silver thiosulfate (STS), water (C) or Promalin (PRO).

| | **STS**[a] | | **C** | | **PRO** | |
|---|---|---|---|---|---|---|
| **Variables** | $R^2$ | $P$ | $R^2$ | $P$ | $R^2$ | $P$ |
| Autumnal bulb mass and leaf area | 0.52 | < **0.001** | 0.26 | < **0.001** | 0.13 | **0.014** |
| Leaf area and bulb mass at final harvest | 0.56 | < **0.001** | 0.89 | < **0.001** | 0.68 | < **0.001** |
| Autumnal bulb mass and mass at final harvest | 0.41 | < **0.001** | 0.25 | < **0.001** | 0.23 | **0.001** |

[a]Degrees of freedom of the error term for STS: 42, C: 50, PRO: 37.

**Supplementary Data 3**: Methods, data (Table S3) and linear regression between leaf glucose concentration analyzed with an HPLC-RID and days since leaf unfolding (Fig. S2).

*Methods*: 500 µL of the supernatant was evaporated under nitrogen, then suspended in 500 µL ultrapure water and filtered through PTFE 0.45 µm filter. Then, 10 µL of this solution was passed through an Aminex HPX-87P 7.8*300 mm, 9 µm chromatographic column at 70 °C for 40 minutes at 0.6 mL $min^{-1}$ in an isocratic gradient. Soluble sugars were detected using a refraction index detector (RID).

*Table S3*: Leaf glucose and fructose concentration per dry mass following days since complete leaf unfolding for Control plants. Sugar concentration was determined by HPLC-RID using the method described above. Each data point represents an individual plant.

| Day since leaf unfolding | Glucose concentration (mg $g^{-1}$) | Fructose concentration (mg $g^{-1}$) |
|---|---|---|
| 6 | 117.2 | 48.4 |
| 6 | 117.0 | 50.2 |
| 8 | 143.4 | 66.2 |
| 13 | 133.9 | 43.9 |
| 13 | 143.3 | 65.0 |
| 20 | 147.5 | 67.6 |
| 21 | 134.1 | 57.0 |
| 21 | 178.4 | 64.9 |
| 21 | 148.4 | 51.1 |
| 23 | 149.5 | 39.7 |
| 29 | 176.5 | 47.4 |

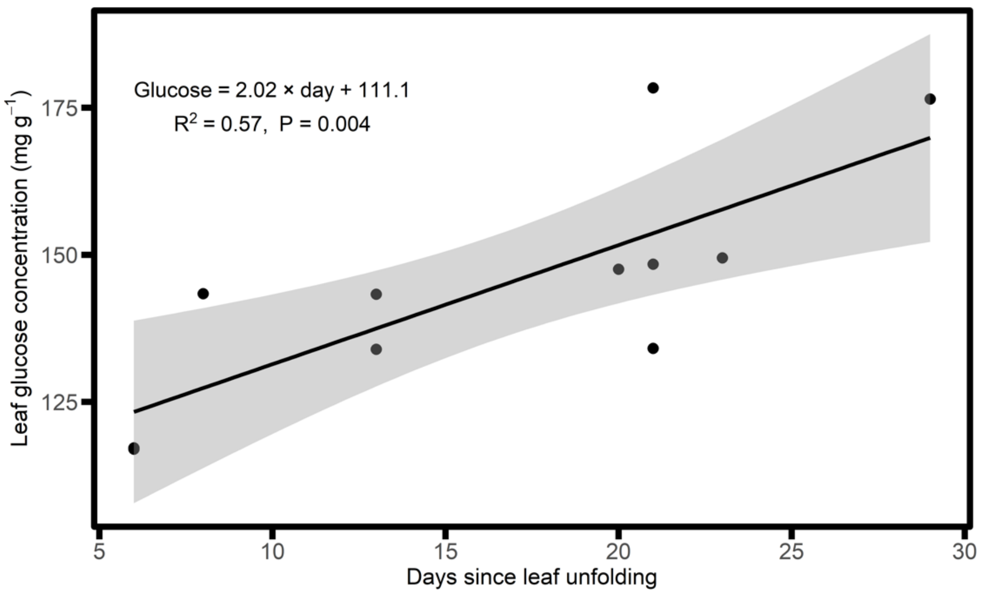


*Fig. S2*: Results of linear regression between leaf glucose concentration and days since complete leaf unfolding for Control plants of *E. americanum*. Glucose concentration was determined by HPLC-RID on plants harvested at maximum photosynthetic rates, at the second foliar spraying, 9 days after the second foliar spraying, and at leaf 5% yellowing.
